# Incorporation of collagen into *Pseudomonas aeruginosa, Staphylococcus aureus*, and *Burkholderia pseudomallei* biofilms enhances their elasticity and resistance against phagocytic clearance

**DOI:** 10.1101/2023.10.25.564018

**Authors:** Xuening Zhou, Ashlee McGovern, Marilyn J. Wells, Deepesh B. Verma, Hailey Currie, Afsana Mimi Raka, Jiachun Shen, Katherine A. Brown, Rae Robertson-Anderson, Vernita D. Gordon

## Abstract

Biofilms are communities of microbes embedded in a matrix of extracellular polymeric substances (EPS) and other components such as proteins. Matrix components can be produced by the microorganisms themselves but can also originate from the environment and then be incorporated into the biofilm. For example, we and our collaborators have recently shown that collagen, a host-produced protein that is abundant in many different infection sites, can be taken up into the matrices of *Pseudomonas aeruginosa* biofilms, altering biofilm mechanics. In an infection, the biofilm matrix protects bacteria from clearance by the immune system, and some of that protection likely arises from the mechanical properties of the biofilm. *P. aeruginosa, Staphylococcus aureus*, and *Burkholderia pseudomallei* are human pathogens notable for forming biofilms *in vitro* and *in vivo* in tissues rich in collagen such as lung and skin. Here, we show that the incorporation of Type I collagen into *P. aeruginosa, S. aureus*, and *B. pseudomallei* biofilms significantly enhances biofilm elasticity and hinders phagocytosis of biofilm bacteria by human neutrophils. Additionally, enzymatic degradation of collagen using collagenase reverses these effects, increasing biofilm susceptibility to neutrophils. Our findings suggest that host materials play significant roles in stabilizing biofilms and may present promising targets for therapeutic interventions.

## Introduction

Biofilms are complex microbial communities that are embedded in a matrix of extracellular polymeric substances (EPS) and other materials, which provides physical and chemical protection against antibiotic treatments and immune clearance [1–7]. Biofilm formation poses a significant clinical challenge as the cause of up to 80% of chronic infections [8], costing the United States healthcare system millions of dollars annually and impacting millions of lives [9, 10]. The most common biofilm-forming bacteria isolated from chronic wounds are *Staphylococcus aureus* and *Pseudomonas aeruginosa* [11]. *Burkholderia pseudomallei* can cause melioidosis, a severe infection that can impact lungs, skin, and other organ systems. *B. pseudomallei* has long been endemic in tropical and subtropical regions of the world, and its regions of endemicity are expanding with climate change [12–14]. For all three of these organisms, their capacity to form biofilms is a major component of their ability to cause persistent diseases that withstand antibiotic treatment and the immune systems. While it is well-known that biofilm-forming microbes can produce EPS, the components of EPS can also originate from the environment. In the context of infections, biofilms can incorporate host materials such as erythrocytes, platelets, fibrin, and DNA [6, 15, 16]. Using an *in vitro* and an *in vivo* experimental model for chronic wounds, we have recently shown that *P. aeruginosa* biofilms incorporate type I collagen into their matrices, resulting in the change of the biofilms’ mechanical properties to be more homogeneous, more relatively elastic, and less compliant [17, 18]. This may arise from collagen adhering to bacteria or entangling physically with the EPS matrix.

Type I collagen is the most abundant protein in all vertebrates [19], including humans. It is the key structural component in major tissues and a mechanical barrier to invasion [20–22]. Type I collagen is commonly found in a wide range of infection sites such as skin, lung, bladder, oropharyngeal, gastrointestinal tracts, and so on [23–28]. Thus, it is plausible that the incorporation of type I collagen into biofilms, and consequent changes in biofilm composition and mechanics, may be widespread well beyond *P. aeruginosa* and chronic wounds specifically. However, the degree to which collagen can be incorporated into non-*Pseudomonas* biofilms and how this impacts the efficacy of the host immune response to the biofilm of any species remains entirely uninvestigated.

Recent research has underscored the potential impact of biofilm mechanics on how effectively the immune system clears infections. Neutrophils are a key component of the human immune system and the most prevalent granulocytes. They phagocytose bacteria and microscopic particles as a way of clearing pathogens. However, the structure of biofilms obstructs phagocytosis physically. Bacterial clusters in biofilms are approximately 100 μm in diameter, while neutrophils are about 10 μm in diameter. For phagocytosis of these rigid, larger targets, neutrophils must detach pieces from the biofilms to engulf them. Our recent work with hydrogel models of biofilm mechanics has revealed that phagocytic success rate and timescale required for successful phagocytosis depend on the elasticity and toughness of the gel target structure [29, 30]. Additionally, we have demonstrated that enzymatic degradation of matrix polymers can significantly alter the mechanical properties of biofilms grown in vitro [31]. Given the increase in relative elasticity and decrease in compliance arising from the incorporation of collagen into biofilms [17, 18], we anticipated that the incorporation of collagen in biofilms should reduce the success of immune cells at phagocytosing biofilm bacteria [32].

*P. aeruginosa* and *S. aureus* are well-known pathogens that form biofilms in a wide range of clinical diseases, including bacteremia, infective endocarditis, and infections of soft tissue and skin [11, 33, 34]. *B. pseudomallei* is largely a lung pathogen, but it can also spread to other parts of the body [35, 36]. Most of the anatomical sites infected by these three organisms are rich in collagen, so bacteria have ample access to this polymer. Similar to most bacteria, *P. aeruginosa, S. aureus,* and *B. pseudomallei* cells are negatively charged at near-neutral pH values [37–40], which should tend to promote electrostatic binding with cationic residues of collagen. Furthermore, these three organisms all have negatively-charged extracellular DNA in their matrices [41–44]; the *S. aureus* biofilm matrix include teichoic acids [45], which have a net negative charge [46]; some strains of *P. aeruginosa* produce copious amounts of negatively-charged alginate [47, 48]. *P. aeruginosa* matrices also contain Psl, which is a net-neutral polysaccharide with charge separation, opening up the possibility of dipole-like attraction with collagen. Extracellular DNA has also been shown to facilitate the adhesion of *B. pseudomallei* during biofilm formation [49]. Taken together, these observations suggest that electrostatics may provide a widespread, generic mechanism for collagen incorporation into biofilms.

Specific interactions of bacteria with collagen are also possible. Proteins bound to the bacterial cell wall of *S. aureus* can adhere to human matrix proteins such as fibrinogen and collagen [50]. For *S. aureus*, adhesion to collagen has been correlated to the development of osteomyelitis and septic arthritis [51]. *B. pseudomallei* produces collagen-like membrane proteins to bind host fibrinogen [52] and also upregulates the expression of host collagenase to reshape the extracellular matrix to their advantage [53, 54].

Thus, collagen may well play important and widespread roles in biofilm infections, but this has been investigated very little. Most investigations of biofilm infections have focused on bacteria-produced materials, and not on materials from the host that may be integrated into the biofilm. Many wound care products in use today aim to break up host material (*e.g.* SANTYL is a collagenase-containing ointment made by Smith & Nephew that is FDA-approved) yet we know little about how this affects biofilm integrity.

Here, for *P. aeruginosa, S. aureus* and *B. pseudomallei* biofilms, we investigate the impact of collagen incorporation on the mechanics, microstructure, and susceptibility to phagocytosis by neutrophils, as well as the effects of targeted enzyme treatments. We found that when biofilms are grown in a collagen-containing medium, neutrophils are less effective at engulfing bacteria and the biofilms exhibit enhanced elasticity; effects that are amplified by increasing the concentration of collagen in the growth medium. To further confirm collagen’s effects on biofilm mechanics and phagocytosis, we treated the biofilms with collagenase, a protease commonly used in wound healing therapy [55, 56]. We found that, for biofilms grown in the presence of collagen, treatment with collagenase increases the phagocytic success rate, and results in less elastic (more dissipative) biofilms. The enhancing effect of collagenase on phagocytic success was greater in cases where the anti-phagocytic effect of collagen was greater. Scanning Electron Microscopy (SEM) imaging of biofilms under all conditions revealed a shield-like “crust” on top of the biofilms that were grown with collagen. This structure disappears following enzyme treatment.

Our findings suggest that incorporating collagen in biofilms has protective effects against host immune reactions. Biofilms grown with collagen form unique microarchitectures, become more elastic, and are more resistant to phagocytosis by human neutrophils. Treatments with collagenase negate these effects and render biofilms vulnerable to phagocytosis by neutrophils. Taken together, our findings suggest that host proteins are likely important components of biofilm infections that contribute to their structural stability and integrity. Although bacteria-produced proteins are widely viewed as contributors to biofilm stability [43, 57–60], the role of host proteins in stabilizing biofilm infections has been neglected. Our work indicates that materials originating in the host should also be considered as potentially important biofilm components. The presence of host proteins in biofilm infections could potentially provide new targets for biofilm dispersion, particularly in cutaneous infections where topical treatments could include the addition of enzymes to degrade biofilm matrix components including host proteins such as collagen.

## Materials and Methods

### Preparation of nutrient media for biofilm growth

Sterile liquid and agar growth media were prepared by dissolving 30 g of LB broth powder (Sigma-Aldrich, Cat.# 154154, NJ, USA) or 30 g of LB agar powder (Sigma-Aldrich, Cat.# 213981, ON, USA), respectively, into 1 L deionized water. Both forms of media were autoclaved on a liquid cycle at 121 °C for 20 minutes. Agar media was allowed to cool, stirring, for 30 minutes at room temperature before being poured into sterile Petri dishes (Fisher Scientific, Cat.# FB0875712, USA) and allowed to solidify overnight, then stored at 4 °C.

### Preparation of collagen (type I collagen) solution

All the pH-balanced collagen solutions in this research were made with refrigerated 4.39 mg/mL Collagen I solution (Collagen I, Rat, Corning 354236, MA, USA). To prepare 2 mL of this solution, 200 µL Phenol Red (Sigma-Aldrich, ACS Cas.# 143-74-8), 200 µL Phosphate-Buffered Saline (PBS, Fisher Scientific, Cat.# BP3991, USA), and 33 µL 1 M NaOH (BDH, ACS Cas.# 1310-73-2) were mixed and slowly added to 1.3 mL of collagen solution. Deionized water was then added to bring the volume to 2 mL, resulting in an orange, viscous liquid, without visible fibers. The solution is stored at 4 °C.

### Growth of biofilms with collagen

A cartoon schematic of the experimental process used to grow biofilms with collagen and measure bacterial engulfment by neutrophils is shown in **Figure 1**.

**Figure 1.**
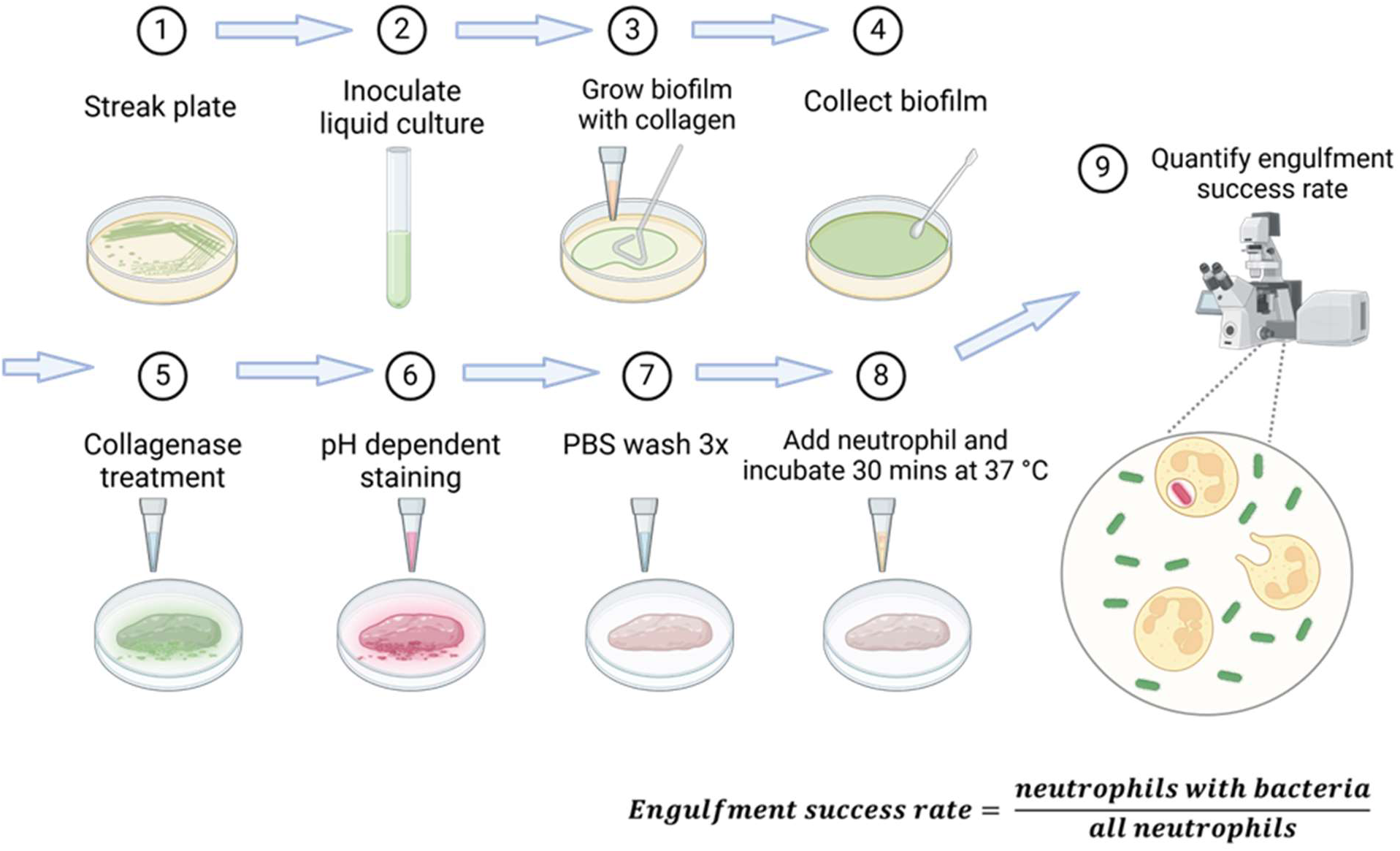
Experimental process used for phagocytosis assays. Biofilms were grown with collagen solution, then collected, stained by pH dependent dye, and then washed multiple times to minimize planktonic bacteria. The biofilms are then incubated with fresh human neutrophils for 30 minutes. After this, neutrophils are extracted and imaged with a confocal microscope to determine the fraction of neutrophils containing bacteria. Created in BioRender. Zhou, X. (2025) https://BioRender.com/u56lxim

The bacterial strains used for biofilm growth in this study were *P. aeruginosa* strains PA01 Wild Type (WT) [61], PA01 Δ*pel* Δ*psl* [62], *S. aureus* strain SA pJY209 [63], and an avirulent *B. pseudomallei* strain Bp82 Δ*purM* [64]. The *P. aeruginosa* strains constitutively express green fluorescent protein (GFP), and the *S. aureus* strain constitutively expresses yellow fluorescent protein (YFP). PA01 Δ*pel* Δ*psl* does not make the extracellular polysaccharides Pel and Psl, which are PA01’s main matrix materials.

The frozen bacterial stock was streaked onto sterile LB agar plates and grown overnight at 37 °C. One colony was selected and cultured in 4 mL of LB broth liquid media and grown shaking overnight at 37 °C. Biofilms were grown by combining 200 µl of overnight culture with 50 µl of 3 mg/mL pH-balanced collagen solution, spread onto LB agar plates with an L-shaped cell spreader (VWR# 220412261B, PA, USA), and incubated statically overnight at 37 °C.

### Treatment of biofilms with collagenase

Solutions of collagenase were prepared by dissolving the enzyme in its powdered form (EMD Millipore Corp., Cat.# 3818971, USA) into phosphate-buffered saline (PBS) at concentrations of 4 mg/mL. Biofilm samples were collected from agar plates gently with a spatula and then transferred to 24-well plates (Fisher Scientific, Cat.# FB012929). Before the phagocytosis assay, each sample was treated with 100 µL of collagenase solution and incubated at 37 °C for 1 hour before staining with pH-dependent dye (see below). All collagenase treatments were of the same dose and duration. For all the control groups, samples were treated with equal volumes of PBS buffer. All other procedures between the control and experimental groups were the same.

### Staining biofilms with pHrodo

A stock solution of pHrodo Red succinimidyl ester (ThermoFisher Scientific, Cat.# P36600, USA) was prepared at a concentration of 1 mg/150 µL in dimethyl sulfoxide (DMSO, Sigma, ACS Cas.# 67-68-5, USA). A working solution was prepared by diluting 1 µL of the stock into 1 mL of PBS. After the enzyme treatment, 500 µL of the working solution were added to each sample and incubated in the dark at room temperature for 30 minutes. After incubation, the samples were washed three times with PBS to remove excess stains and planktonic bacteria.

### Isolation of human neutrophils

Work with neutrophils was approved by the Institutional Review Board at the University of Texas at Austin (Austin, TX) as Protocol No. 2021-00170.

Human neutrophils from adult volunteer blood donors were isolated following a previously published protocol [29, 30]. In brief, freshly drawn whole blood was mixed with an equal volume of a solution containing 3% dextran (Sigma-Aldrich, ACS Cas.# 9004-54-2, Denmark) and 1.8% sodium chloride (Fisher Scientific, ACS Cas.# 7647-14-6). After 20 minutes undisturbed, most red blood cells formed aggregates and dropped to the bottom of the tube. The supernatant was centrifuged (Eppendorf 5810R, A-4-62 Rotor) for 10 minutes at 500 x g to separate the remaining blood cells from the plasma. The supernatant, mainly containing plasma, was discarded, and the remaining pellet was suspended in 10 mL Hanks Buffered Salt Solution (HBSS, Gibco Laboratories, Cat.# 14175095, Frederick, USA) without calcium or magnesium. This suspension was slowly added to the top of 4 mL Ficoll-Paque density gradient (GE Healthcare #17-1440-02, Uppsala, Sweden) and centrifuged for 40 minutes at 400 x *g*. The pellet was mixed with 4 mL of deionized water for 30 seconds to lyse the residual red blood cells, then stabilized by mixing with 4 mL of filter-sterilized 1.8% NaCl solution. Following 5 minutes of centrifugation at 500 x *g*, a white pellet of neutrophils was resuspended in 800 µL HBSS with calcium and magnesium and 200 µL human serum. Each lithium-heparin-coated tube of blood drawn yielded 1 mL of neutrophil-only solution. In experiments where more than one tube of blood was drawn, the resulting neutrophil solutions were thoroughly mixed into one tube by pipette. Isolated neutrophils were quantified by microscopy for consistency.

### Phagocytosis assay

This assay was done following our previously-published protocol [65]. Briefly, each biofilm sample was mixed with 200 µL of the isolated neutrophil suspension in the 24-well plate and incubated at 37 °C for 30 minutes. The supernatant in the wells was discarded by pipetting. The pellet at the bottom of each well was washed with 200 µL of HBSS with calcium and magnesium, then transferred and resuspended in a centrifuge tube. The tube was spun down for 3 minutes with a Mini centrifuge (Bioexpress, Max. speed 6,000 rpm). 150 µL of the resulting supernatant were removed, and the pellet was resuspended with the remaining supernatant. The resuspended liquid was transferred to an image spacer on a slide (Fisher Scientific, Cat.# 12-544-1, PA, USA) and covered with a coverslip (Fisher Scientific 12542B, PA, USA). The sample was then observed using confocal laser scanning microscopy (Olympus IX71 inverted confocal microscope, 60X oil-immersion objective), and analyzed with FluoView FV10-AWS version 04.02 from Olympus America. The fluorescence was visualized by overlaying bright-field image, red, and green fluorescence channels. Red fluorescence within a neutrophil indicates that one or more bacteria have been engulfed in the highly acidic phagosome (Fig. S1). The phagocytic success rate was defined as the percent of neutrophils in a sample that contained at least one engulfed, red bacterium. At least 100 neutrophils were counted per sample and assessed for phagocytic success. The numbers of biological replicates and total cells counted are shown below in Table 1. Neutrophils from three different volunteer donors were used.

**Table 1.** Numbers of biological replicates and neutrophils counted for phagocytosis assays on biofilms.

| Bacterial Strain | Growth and treatment conditions | WT PA01 | PA01 $\Delta pel\Delta psl$ | SA pJY209 | Bp82 |
| --- | --- | --- | --- | --- | --- |
| Number of biological replicates |  | 7 | 7 | 7 | 6 |
| Total neutrophils counted | 0% collagen, untreated | 1162 | 1074 | 1072 | 971 |
|  | 0% collagen, + collagenase | 1037 | 1195 | 1028 | 1100 |
|  | 10% collagen, untreated | 1232 | 1150 | 1176 | 991 |
|  | 10% collagen, + collagenase | 1070 | 963 | 1262 | 967 |
|  | 20% collagen, untreated | 977 | 1288 | 1054 | 1005 |
|  | 20% collagen, + collagenase | 1142 | 1336 | 956 | 1057 |

### Data Analysis

Statistical significance was determined by Student *t*-tests and single-factor ANOVA tests in Microsoft Excel. The threshold for significance was set for *p*-values less than or equal to 0.05. The *p*-values for each dataset and statistical test are shown in Table 2.

**Table 2.**
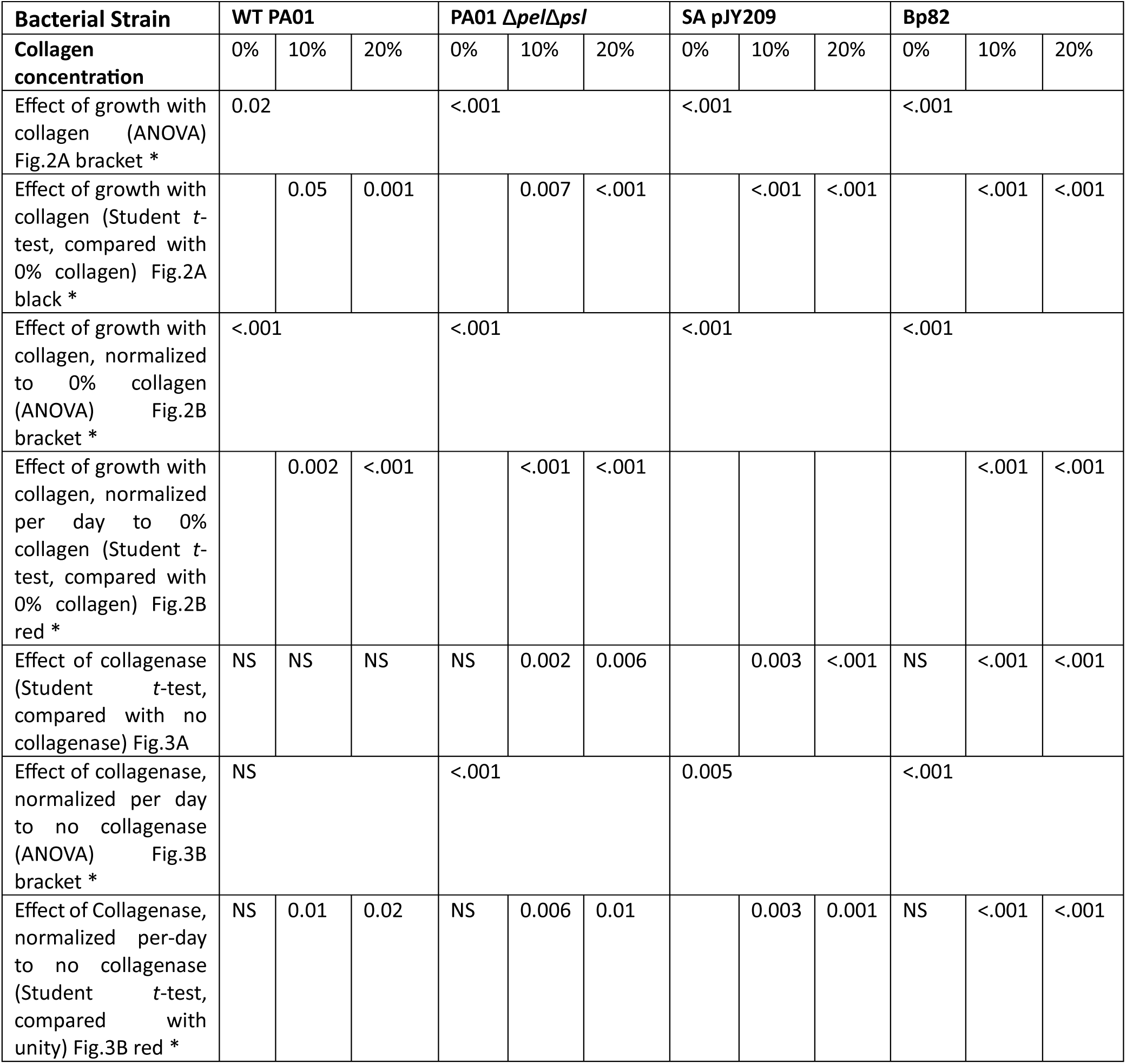
p-values for statistical tests to assess the impact, on phagocytic success, of growth with collagen and treatment with collagenase. The type of test used (ANOVA or Student t-test) is indicated for each row.

### Preparation of alginate hydrogels

Alginate gels were prepared by dissolving 4% sodium alginate (Sigma-Aldrich, Cat.# 180947) and 1% Bovine Serum Albumin (HyClone, GE Life Sciences Cat# SH30574.01) in deionized water. This solution was stirred constantly on a hot plate for at least one hour. Then, the solution was transferred to 24-well plates and mixed with fluorescent tracer beads (polystyrene Dragon Green beads, diameters 0.955 mm, Cat# FSDG004; Bangs Laboratories, Fishers, IN, USA) at a concentration of 1 µL tracer per 100 µL of alginate solution. The mixture is stored at 4 °C overnight to form hydrogel samples. Information on replicate numbers and statistical outcomes for these experiments are given in Table 3.

**Table 3.** Statistics for phagocytosis assays on hydrogels and bacterial suspensions.

|  | Alg. Untreated | Alg. + collagenase | *P. aeruginosa* untreated | *P. aeruginosa* + collagenase | *E. coli* untreated | *E. coli* + collagenase |
| --- | --- | --- | --- | --- | --- | --- |
| Biological replicates | 3 | 3 | 3 | 3 | 3 | 3 |
| Technical replicates | 2 for each biological replicate |  |  |  |  |  |
| Total neutrophils counted | 847 | 887 | 826 | 788 | 782 | 733 |
| Student *t*-test *p*-value | NS |  | NS |  | NS |  |
| ANOVA *p*-value | NS |  | NS |  | NS |  |

### Preparation of bacterial suspension

We used the DH5α strain of *Escherichia coli* and previously mentioned PA01 WT strain. The frozen stock was streaked onto sterile LB agar plates and grown overnight at 37 °C. One colony was selected and cultured in 4 mL of LB liquid medium, shaking overnight at 37 °C. The following day, the liquid culture was centrifugated at 10,000 x *g*, resuspended with 100 µL collagenase solution or PBS, and incubated for 1 hour. Then 500 µL of the pHrodo working solution (see *Staining biofilms with pHrodo*) were added to each sample and incubated at room temperature for 30 minutes. The stained samples were washed 3 times with PBS by centrifuging at 10,000 x *g* to remove the extra stain, resuspended in PBS, and diluted to an optical density (OD600) of 1. After the dilution, 100 μL of the bacterial suspension were added to each well in a 24-well plate before the *phagocytosis assay* (described above). Information on replicate numbers and statistical outcomes for these experiments are given in Table 3. As an initial study and for internal consistency within data sets, all planktonic *P. aeruginosa* and *Escherichia coli* experiments were performed with blood from 3 volunteers.

### Particle tracking microrheology

To determine the rheological properties of the biofilms, we track the thermal motion of tracer polystyrene microspheres (1 µm, carboxylate-modified, red fluorescent, polystyrene latex, from Bangs Laboratories, Fishers, IN) embedded in biofilms. Overnight liquid culture was mixed with microspheres (125:1) and thoroughly resuspended by pipetting, then incubated overnight at 37 °C in glass-bottom dishes (No. 1 Uncoated, Matsunami Glass) to form a transparent, gel-like biofilm. The 1 µm size and carboxylate modification allow the particles to embed into the biofilm matrix as it develops and also satisfy microrheology requirements of being larger than the matrix mesh size and non-interacting with the matrix materials (demonstrated in [66, 67]). These biofilms are grown under different conditions than those for engulfment experiments, and we expect biofilm properties to be dependent on growth conditions [68]. To visualize tracer motion, we record 8-second videos of diffusing tracers at a frame rate of 33 fps using an Olympus IX71 phase contrast microscope, 60X objective, and an Orca Flash4.0v2 camera. Each of the 264 frames of each video are 2048 x 2048 square-pixels that correspond to 157.5 x 157.5 µm^2^. For each condition, we captured 3-10 videos per replicate for at least three biological replicates. The number of tracks reported for each sample varies and is listed accordingly in Table 4.

**Table 4.**
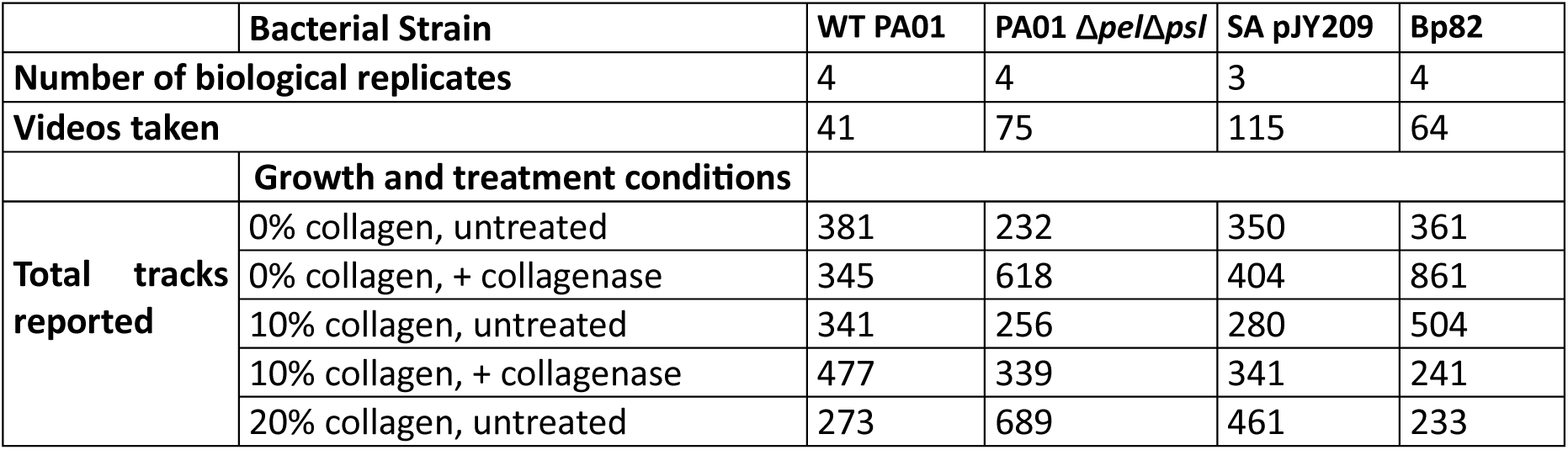
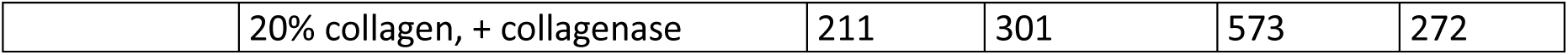
Numbers of tracks reported for microrheology assays on biofilms.

We use the particle-tracking plug-in ‘Trackmate’ in Fiji [69] to determine the time-dependent positions (*x*(*t*), *y*(*t*)) of each particle in the video (Table 4). From these trajectories we use trackpy [70] to determine the mean-squared displacement (*MSD*(Δ*t*) = 〈(Δ*x*(Δ*t*))^2^ 〉 + 〈(Δ*y*(Δ*t*))^2^ 〉 as a function of lag time Δ*t* (**SI Fig S2-S6**).

We determine error bars on MSDs using the following procedure for each combination of bacterial strain, collagen concentration, and collagenase treatment (or not) tested: For each lag time, the standard deviation of the MSD over all trajectories measured was computed and divided by the square root of the number of biological replicates. **Supplementary Figure S6** shows the weighted averages of the MSDs for each biofilm condition and lag time, where the weight is determined by the (number of trajectories in that technical replicate / total number of trajectories).

For normal thermal Brownian motion, the MSD scales linearly with lag time, and the constant of proportionality provides a measure of the diffusion coefficient *D*:

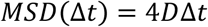

In environments that restrict or confine particle displacements, such as viscoelastic and crowded systems [71–74], the scaling of MSD with lag time is typically sublinear, a phenomenon known as anomalous subdiffusion, and described as

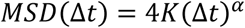

where *α* < 1 is the anomalous scaling exponent and *K* is the generalized transport coefficient [71, 75]. Lower *α* and *K* values indicate more restricted and slower motion. Error bars in *α* and *K* values are determined by using the same procedure described above for determining error bars on MSDs.

To directly determine the viscoelastic properties from the MSDs, we use the generalized Stokes−Einstein relation (GSER) to extract the frequency-dependent elastic and viscous modulus, *G*′(*ω*) and *G*″(*ω*), which are measures of how elastic or solid-like (*G*′) and viscous or liquid-like (*G*″) a material is [71, 76, 77]. *G*′ and *G*″ values reported below are the average over the low-frequency values (*X* < *ω* < *X*) where the values are relatively independent of *ω*.

### Scanning Electron Microscopy Imaging of Biofilm Structure

Biofilms were grown overnight on 50×50mm Aclar (Ted Pella, Redding, CA) substrate placed in 24-well plates (Corning, Corning, NY) containing 1 mL LB media. The samples were then fixed for microscopy following our published enhanced biofilm fixation methods [78]. In summary, the biofilms were fixed overnight using 4% glutaraldehyde (Electron Microscopy Sciences, Hatfield, PA), 2% paraformaldehyde (Electron Microscopy Sciences, Hatfield, PA), and 0.15% Alcian blue (Sigma, St. Louis, MO) in 0.1 M sodium cacodylate buffer at pH 7.4 (Electron Microscopy Sciences, Hatfield, PA). The samples were washed three times with 0.1 M sodium cacodylate buffer and then post-fixed in a solution of 1% osmium tetroxide (Ted Pella, Redding, CA) and 1% tannic acid (Electron Microscopy Sciences, Hatfield, PA). After 2 hours, the samples were gradually dehydrated using 30%, 60%, 75%, 90%, 95% and absolute ethanols (Thermo Fisher, Waltham, MA) and hexamethyldisilazane (HMDS, Ted Pella, Redding, CA), then coated with 12 nm platinum/palladium using a Cressington 208HR sputter coater (Ted Pella, Redding, CA). SEM imaging (Zeiss Supra field emission) was conducted with an SE2 detector and a 5 kV accelerating voltage.

## Results and Discussion

### The incorporation of collagen in biofilms impedes the engulfment of bacteria by neutrophils

We grew biofilms from *P. aeruginosa* PA01 wild type (WT), PA01 Δ*pel*Δ*psl, S. aureus* SA pJY209, and *B. pseudomallei* Bp82 in LB media, in the presence of three concentrations of collagen: 0%, 10%, and 20%. The biofilms were stained with pH-dependent dye and then incubated with freshly isolated human neutrophils for 30 minutes. Imaging with confocal laser fluorescence scanning microscopy allows us to identify bacteria that have been internalized into neutrophils’ highly acidic phagosomes because they fluoresce in a different color from external bacteria, including those stuck to the surfaces of neutrophils (**Fig. S1**). For each sample, imaged fields of view were chosen at random until the number of analyzed neutrophils exceeded 100. Usually, 3-10 fields of view were analyzed for each sample, with each field of view containing 1-120 neutrophils. Phagocytic success is quantified by measuring the percentage of imaged neutrophils that contain bacteria. Representative images for each bacterial strain and collagen concentration are shown in **Supplementary Figures S7-S10**.

In the absence of collagen, the average phagocytic success rate of neutrophils applied to biofilms of WT PA01 *P. aeruginosa* was 55%. However, this figure decreased to 40% and 31% when the biofilms were grown with 10% and 20% collagen, respectively (**Fig. 2A**). For neutrophils applied to biofilms of *P. aeruginosa* Δ*pel*Δ*psl* (which does not produce the two matrix polymers that are most important for PA01), the average phagocytic success rate declined from 51% (when biofilms were grown without collagen) to 35% and 24% when biofilms were grown in the presence of 10% and 20% collagen, respectively (**Fig. 2A**). For neutrophils applied to biofilms of *S. aureus*, the phagocytic success rate also declined from 65% (when biofilms were grown without collagen) to 38% and then to 27% with increasing collagen concentration (**Fig. 2A**). For neutrophils applied to biofilms of *B. pseudomallei* Bp82, the phagocytic success rate also declined from 22% (when biofilms were grown without collagen) to 14% and then to 11% with increasing collagen concentration (**Fig. 2A**). All of these results are statistically significant.

**Fig 2.**
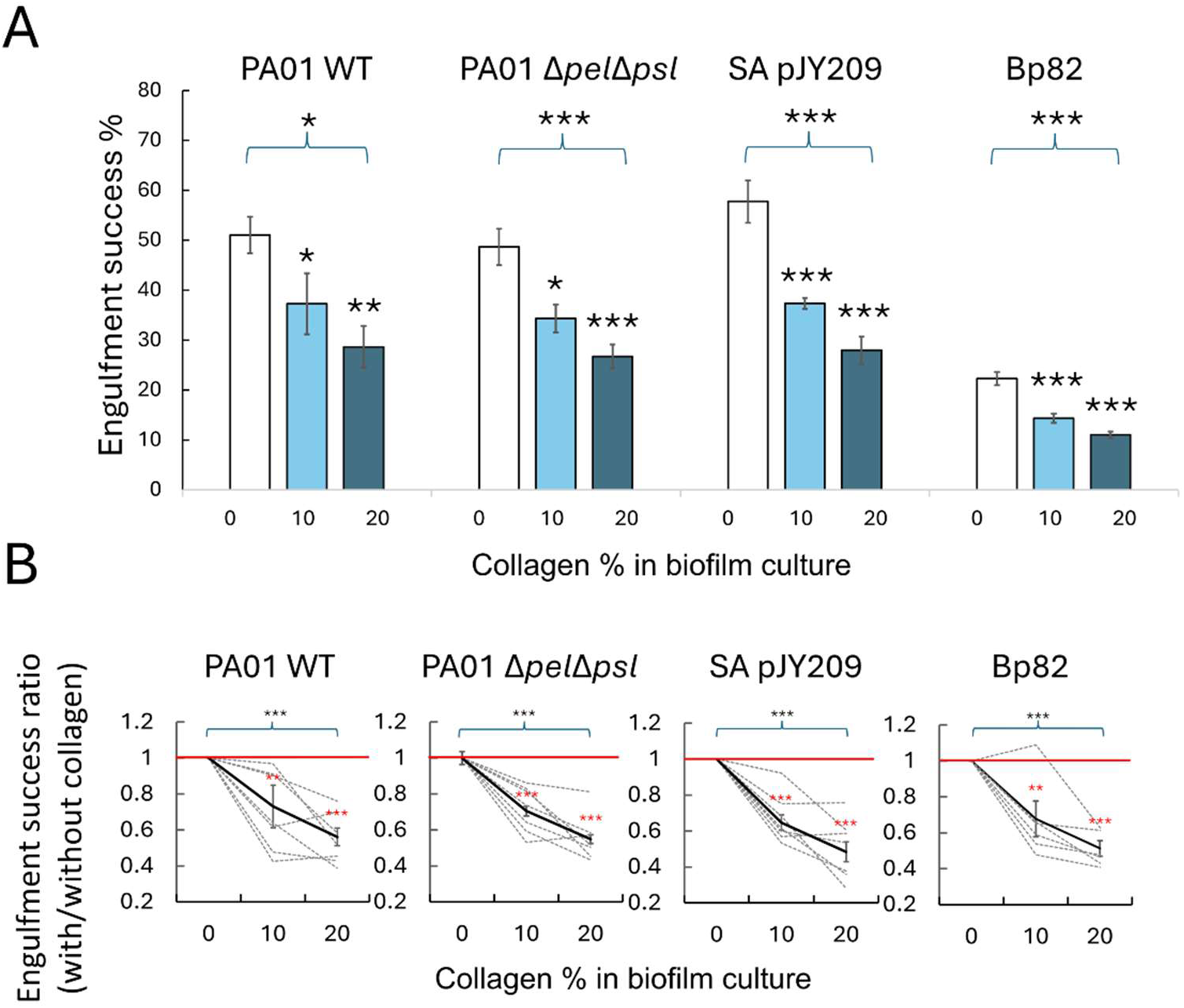
Collagen impedes engulfment of biofilm bacteria by neutrophils. (A) For neutrophils applied to biofilms, the phagocytic success rates are negatively correlated with the concentration of collagen in which the biofilms were grown. This is true for biofilms grown from four different bacterial strains of three different species. Statistical markers over individual bars indicate a comparison to the corresponding control (0% collagen for each bacterial strain). Statistical markers with brackets indicate overall comparison within a strain, for all three collagen concentrations. (B) Collagen’s effect on engulfment success, shown as fold changes from the control condition (0% collagen for each bacterial strain). Dashed lines represent individual biological replicates, grown and measured in parallel on the same day. Solid lines represent the average of replicates across different days. The red horizontal lines indicate unity, for visual comparison with measured values. The error bars represent the standard error of the mean. Statistical markers in red indicate comparing the fold change to unity (which would mean no change), using Student t-tests. Statistical markers with brackets indicate overall comparison within a strain, for all three collagen concentrations, using the ANOVA test. (A and B) Statistically significant differences are shown by * for *p* < 0.05, ** for *p* < 0.01, and *** for *p* < 0.001. The p-values and replicate numbers (N) for each strain are shown in Tables 1 and 2, respectively.

Our results suggest that in three species of clinically important biofilm-forming bacteria, the incorporation of collagen into biofilms inhibits phagocytosis by neutrophils. In total, four bacteria strains producing different EPS components were tested [17, 79, 80]. The trend of decreasing phagocytic success with increasing collagen concentration is seen for all four strains tested.

To quantify the effects on phagocytic success upon the addition of 10% and 20% collagen, we measured the fold change, comparing to the success rate at 0% collagen (**Fig. 2B**). Here, a fold change of unity represents no change. In contrast, a fold change greater or less than unity represents an increase or decrease, respectively, in phagocytic success. This normalization approach is intended to account for day-to-day variations in both biofilm properties and neutrophil efficacy, and it parallels day-by-day normalization approaches we have used in previous work [30, 81, 82]. In most cases, the sizes of day-to-day fluctuations in engulfment success are comparable to the sizes of average changes in engulfment success. In **Figure 2B**, each dotted line represents the fold changes measured on one day from one biological replicate, while solid lines represent the average of fold changes measured for different biological replicates on different days.

Adding 10% collagen decreased the engulfment success rate of *P. aeruginosa* WT strain to 0.7 of its original value, Δ*pel*Δ*psl* strain to 0.7 of its original value, *S. aureus* pJY209 strain to 0.6 of its original value, and *B. pseudomallei* Bp82 to 0.6 of its original value. With 20% collagen, those fold changes became 0.6, 0.5, 0.5, and 0.5, respectively. PA01 WT biofilms contain more EPS, in amount and variety, than do Δ*pel*Δ*psl* biofilms. This may be related to why the incorporation of collagen, another polymer, has more impact on the success of neutrophils at engulfing bacteria from Δ*pel*Δ*psl* biofilms than from WT. The ability of *S. aureus* to specifically bind to collagen may be related to why the incorporation of collagen has more impact on the success of engulfing bacteria from *S. aureus* biofilms than from either *P. aeruginosa* strain [50]. Although the interaction between *B. pseudomallei* Bp82 cells and collagen remains unclear, this interaction might play a role in *B. pseudomallei*’s dissemination to human tissues and organs in acute infections. *B. pseudomallei* not only produces bacterial collagenases [83] but also augments the expression of matrix metalloproteinases by infected human skin fibroblasts [53]. These enzymes digest collagen and other ECM components in human tissues [84], and induce cellular changes during inflammation that facilitate bacterial immune invasion [85], thereby causing severe tissue damage and supporting the spread of bacteria into the circulatory system. Therefore, understanding how *B. pseudomallei* interacts with collagen and collagenase is vital for understanding the pathogenesis of this pathogen.

### Collagenase treatments of biofilms promote neutrophil engulfment

We have previously demonstrated that enzymes that target alginate and extracellular DNA, when these are the main polymer components of the *P. aeruginosa* biofilm matrix, can compromise the biofilm’s mechanical properties [31]. We have also previously shown a potential connection between bacterial biofilm mechanics and phagocytic success [29, 30] and that incorporating collagen into *P. aeruginosa* biofilms increases their relative elasticity and decreases their creep compliance [17, 18]. Therefore, we expect that enzymatically breaking down collagen within biofilms might enhance the success rate of phagocytosis of biofilm bacteria by neutrophils. To investigate this, we grew biofilms with different concentrations of collagen (0%, 10%, and 20%) and then treated them with collagenase before incubating them with neutrophils. We then measured phagocytic success as above. Representative images for each bacterial strain and collagen concentration are shown in **Fig. S3-S6**. For the control groups, an equal volume of PBS was used instead of collagenase solution. Each pair of test and control experiments were done on the same day, on the same biological biofilm, using neutrophils isolated on that day in the same batch from the same donor.

For biofilms without collagen, collagenase treatment had no significant effect on the average phagocytosis success rate. For PA01 WT biofilms grown with 10% and 20% collagen, collagenase treatment increased by 9% and 8%, compared to their respective untreated control; however, these increases were not statistically significant (**Fig. 3A**). For PA01 Δ*pel*Δ*psl* biofilms grown with 10% and 20% collagen, collagenase treatment increased the average phagocytosis success rate by 17% and 18%, respectively (**Fig. 3A**). For *S. aureus* pJY209, treatment with collagenase increased the average phagocytic success rate by 21% for biofilms grown with 10% and 20% collagen (**Fig. 3A**). For *B. pseudomallei* Bp82, treatment with collagenase increased the average phagocytic success rate by 20% and 43% for biofilms grown with 10% and 20% collagen, respectively (**Fig. 3A**). For PA01 Δ*pel*Δ*psl, S. aureus,* and *B. pseudomallei* biofilms, the increases in engulfment success upon collagenase treatment were all statistically significant.

**Fig 3.**
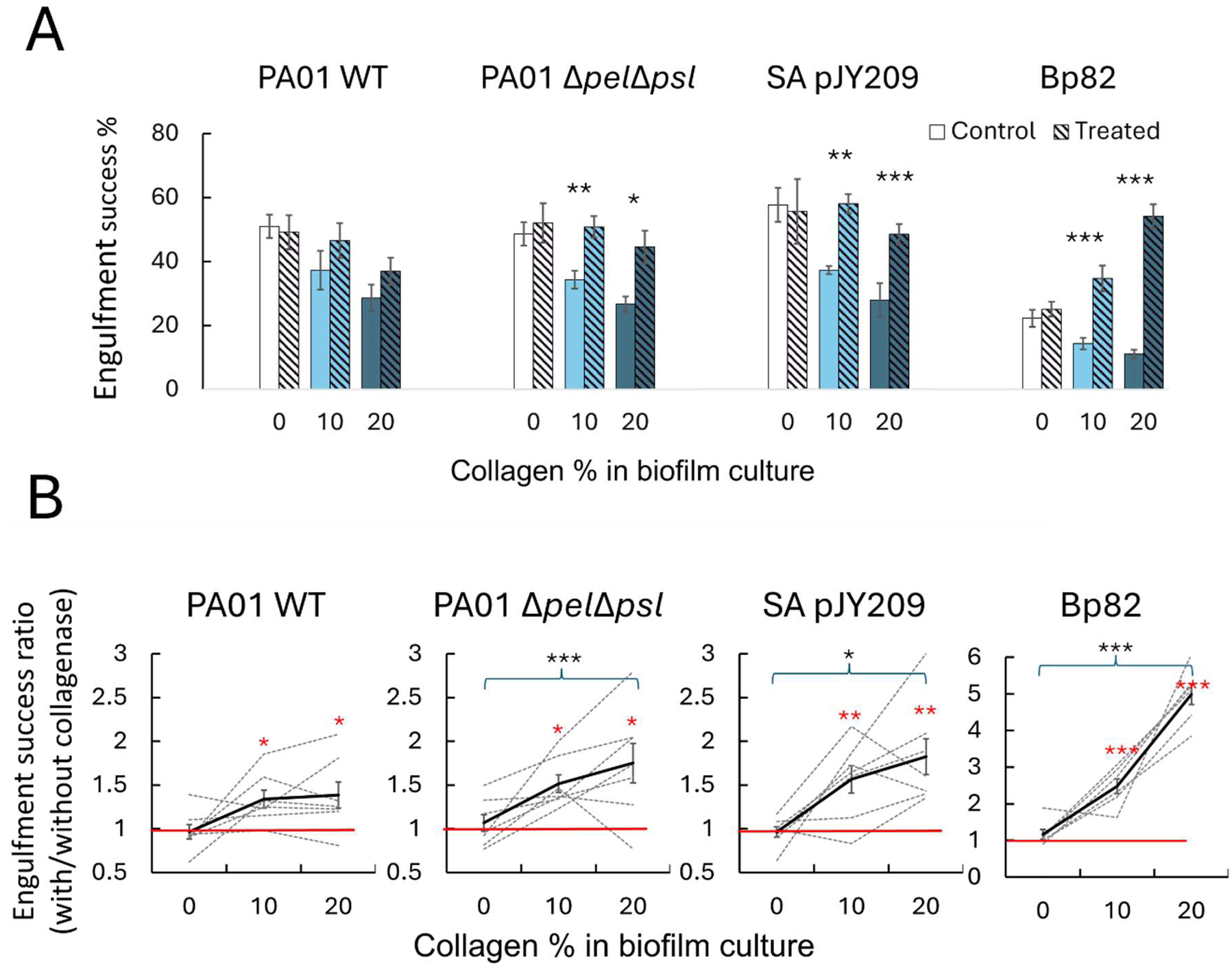
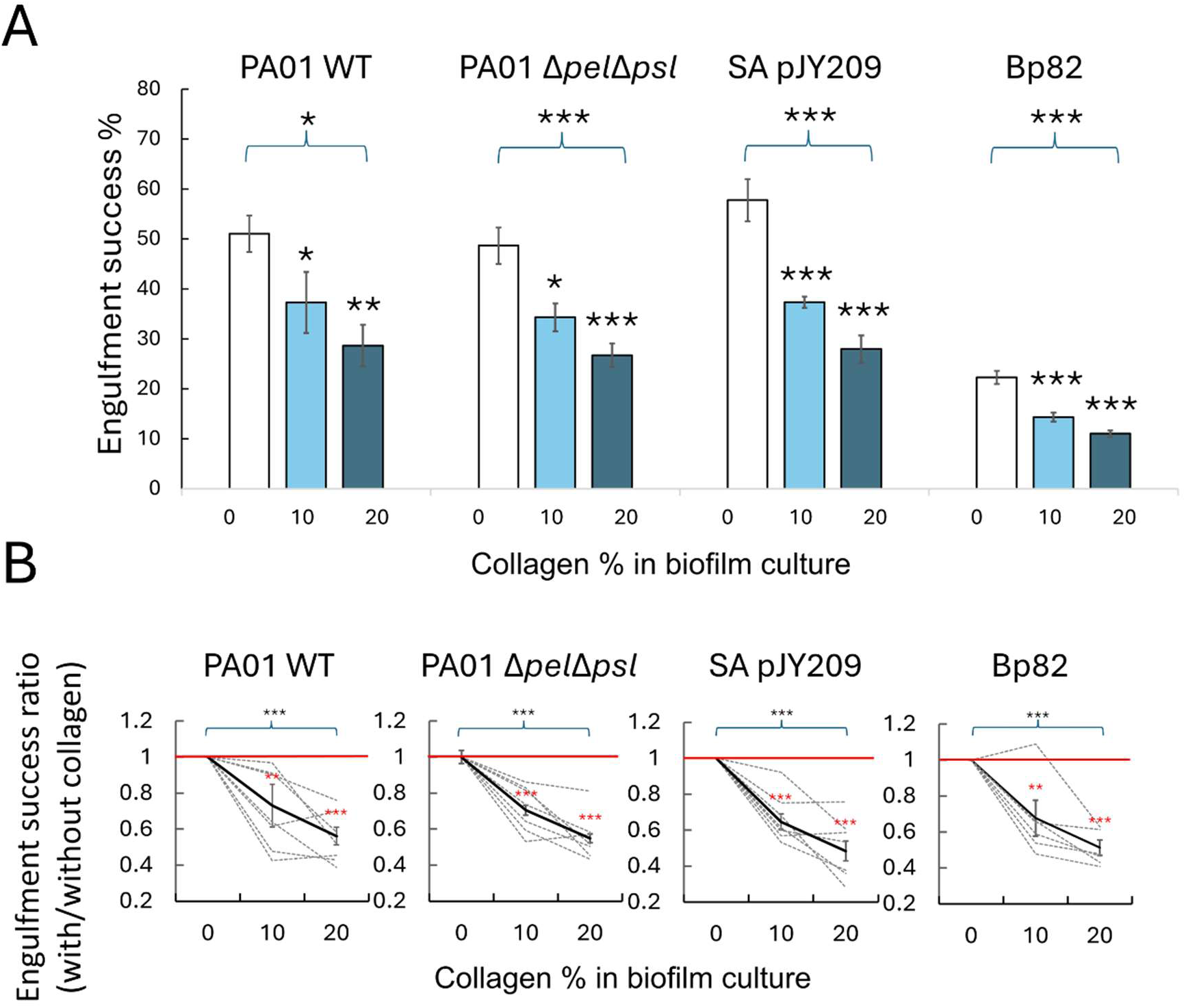
When biofilms grown with collagen are treated with collagenase, the subsequent phagocytic success of neutrophils increased. (A) Engulfment success rates for untreated (control, solid bars) biofilms paired with biofilms that were treated with collagenase (hashed bars) before incubation with neutrophils. Significance markers indicate a statistically-significant difference between the treated and control biofilm, using the Student t-test. (B) Collagenase’s effect on engulfment success, shown as the fold change from the untreated (control, no collagenase) case for each condition. Dashed lines represent individual biological replicates, grown and measured in parallel on the same day. Solid black lines represent the average of replicates across different days. The red horizontal lines indicate unity, for visual comparison with measured values. Above the data points, red statistical markers indicate a comparison to unity using the Student t-test. Black statistical markers with brackets indicate comparison of the three sets of values using ANOVA. (A and B) Statistically significant differences are shown by * for p < 0.05, ** for p < 0.01, and *** for p < 0.001. The p-value between 2 groups came from the student t-test, and the overall p-value for each strain came from the ANOVA test. Error bars represent the standard error of the mean. The p-values and replicate numbers (N) for each strain are shown in Tables 1 and 2, respectively.

We quantified the impact of collagenase treatment on engulfment success by calculating the fold change of the phagocytic success rate for each biological replicate at each collagen concentration, compared with the same biological replicate and collagen concentration sham-treated with PBS instead of collagenase (**Fig. 3B**). A fold change of unity indicates no change, while a fold change greater (or less than) unity indicates an increase (or decrease) in phagocytic success. This approach normalizes the data to account for day-to-day fluctuations in the fold change associated with collagenase treatment. In many cases, the sizes of day-to-day fluctuations in the fold change are comparable to the sizes of the average fold changes; this can be seen by comparing dashed lines in **Fig. 3B**, which each represent the fold change measured on one day for one biological replicate, with solid lines in **Fig. 3B**, which represent the average of fold changes measured for different biological replicates on different days.

For biofilms grown with 10% collagen, collagenase treatment increased the engulfment success rate of *P. aeruginosa* WT strain to 1.3 of its original value, Δ*pel*Δ*psl* strain to 1.5 of its original value, *S. aureus* pJY209 strain to 1.6 of its original value, and *B. pseudomallei* Bp82 to 2.5 of its original value. For biofilms grown with 20% collagen, those fold changes due to collagenase treatment became 1.4, 1.8, 1.8, and 5.0, respectively. It should be noted that although the raw data for PA01 WT (**Fig. 3A**) did not show statistical significance, normalization of the day-to-day variation resulted in statistical significance, determined using student t test (**Fig. 3B**). For each bacterial strain tested, collagenase treatment of biofilms grown with collagen resulted in an increased phagocytic success rate, with statistical significance (**Fig. 3B**).

### Outside the biofilm context, collagenase does not impact phagocytic success

Our data showed that for biofilms lacking collagen there were no statistically significant changes in phagocytic success induced by treatment with collagenase. These results align with our expectations, as collagenase does not break down the EPS in these biofilms. However, prior studies have shown that collagenase can modify and regulate the bioactivity of chemokines upstream of Interleukin-8, thus possibly involved in the recruitment and activation of neutrophils [86, 87]. We did two sets of experiments to evaluate the impact of collagenase on the neutrophil activity in our experimental conditions, as follows:

To assess the effect of collagenase on neutrophils’ extraction of sub-portions of a large target structure, we performed engulfment assays using alginate hydrogels that contained no collagen. Tracer beads that were incorporated into the alginate solution before it gelled were used to measure phagocytic success; this is the same approach we used in our earlier work [29, 30]. The hydrogels were subjected to collagenase treatment of the same dose as in the biofilm experiments and then incubated with fresh human neutrophils for 1 hour. Representative images are shown in **Fig. S11**. In this case, there was no statistically significant effect of collagenase treatment on the engulfment success rate (**Fig. S12** and Table 3). This parallels our finding that, for biofilms grown without collagen, collagenase has no statistically significant effect on phagocytic success (**Fig. 3**).

Further, to assess whether collagenase affects the interaction between neutrophils and live bacteria, engulfment experiments were conducted using planktonic suspensions of *P. aeruginosa* and *E. coli* [88]. Mid-log phase cultures were washed multiple times by centrifugation to minimize the remaining EPS in the suspension, then treated with collagenase of the same dose as above, then stained with pH-dependent dye, washed again by centrifugation to remove the excess dye, and incubated with neutrophils for one hour. The control groups were treated with equal volumes of PBS instead of collagenase. Representative images are shown in **Fig. S11**. The phagocytic success rate was evaluated as above. The results revealed that the addition of collagenase did not yield statistically significant changes in the engulfment success rate for planktonic bacterial cells (**Fig. S12** and Table 3).

Thus, in collagen- and EPS-free systems, we do not find a statistically significant enhancement of phagocytic success by collagenase. Therefore, we conclude that collagenase’s effect on enhancing the engulfment of biofilm bacteria primarily arises from the enzymatic digestion of collagen incorporated into the biofilm matrix rather than any direct biochemical influence on the neutrophils.

### Incorporated collagen suppresses particle transport in biofilms

Biofilms are viscoelastic materials, meaning that they exhibit a combination of both solid-like and fluid-like responses to mechanical stresses and strains, depending primarily on the composition and interaction of its EPS polymers. This viscoelastic nature can lead to anomalous and restricted transport of particles through the biofilm [89, 90] that may play a key role in protection against mechanical or chemical challenges such as attack by neutrophils and antibiotic treatment [32, 91–93].

To analyze how growth in the presence of collagen impacts the transport of particles through biofilms, we performed particle tracking microrheology, as described in Methods. Briefly, we evaluated the mean-squared displacements (MSDs) of particles embedded in biofilms (**Fig S2-S6**), from which we determined the anomalous scaling exponent *α* and transport coefficient *K* by fitting the data to *MSD* = 4*K*(Δ*t*)^a^. In a purely viscous fluid, particles undergo normal Brownian diffusion, in which *α* = 1 and *K* equates to the diffusion coefficient, with larger *K* values indicating faster motion. In viscoelastic environments, or otherwise confined or crowded systems, particles often exhibit anomalous subdiffusion, characterized by *α* < 1, with lower *α* values indicating more strongly restricted motion. Higher *K* values still indicate faster motion, but the units differ from that of normal diffusion coefficients and depend on *α*.

For biofilms of *P. aeruginosa* PA01 wild type (WT), PA01 Δ*pel*Δ*psl, S. aureus* SA pJY209, and *B. pseudomallei* Bp82, growth with collagen resulted in significantly suppressed particle motion compared to the 0% collagen cases, as seen by the MSDs generally shifting to lower values (lower *K*) and displaying more shallow slopes (lower *α*) (**Fig S6**).

The qualitative trends seen in the MSDs are quantified in **Fig 4**. As shown, collagen generally reduced the transport coefficients in biofilms, with SApJ209 and Bp82 displaying the most significant decreases in *K* values (**Fig 4A**). Moreover, all untreated collagen cases displayed subdiffusive behavior, i.e. *α* < 1, with exponents that were generally lower than those for 0% collagen cases (**Fig 4B**). Upon collagenase treatment, the transport coefficients and exponents for nearly all biofilms and collagen percentages increased, with Bp82 displaying the largest effect (**Fig 4**). The impact of collagen on transport properties suggests that growth in collagen results in a significantly more confined and viscoelastic environment that hinders the motion of small particles. This suppressed transport may play a role in the reduced ability of neutrophils to engulf micron-sized particles, as we discuss further below.

**Fig 4.**
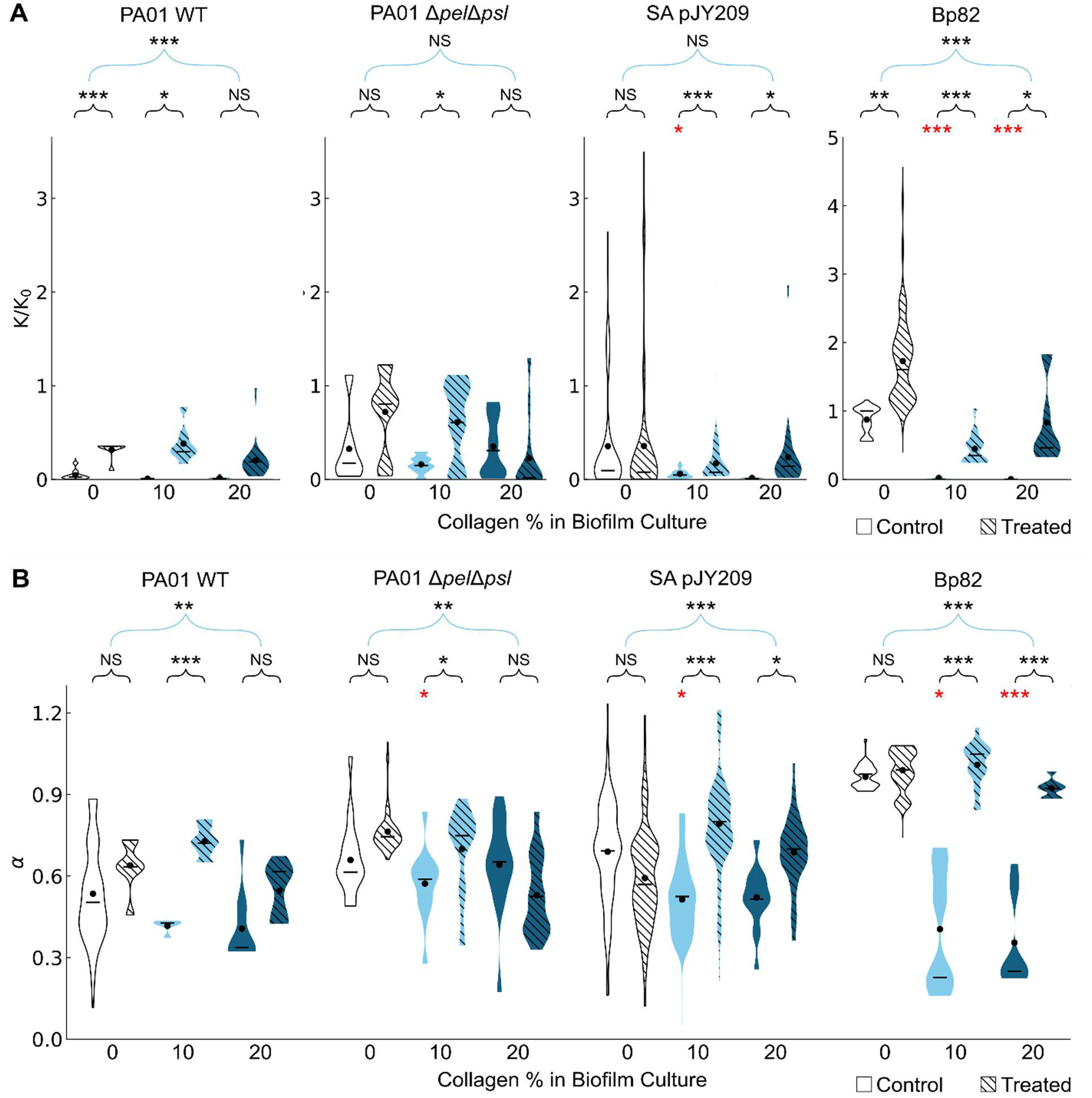
Diffusion and transport of particles within biofilms both with and without the presence of collagenase. (A) Transport coefficient normalized by transport in water *K*/*K*_O_ for the untreated biofilms (control, solid violin plots) and collagenase-treated (hashed violin plots) biofilm cases. (B) Anomalous scaling exponent *α* for untreated (solid)) and collagenase-treated (hashed) biofilms. The black dot is the weighted average, and the line is the median. (B) (A and B) Black statistical markers with blue brackets indicate results of the comparison of the six sets of values (all three collagen concentrations, without and with collagenase treatment) using ANOVA. Black statistical markers with black brackets indicate the results of the comparison of the biofilm grown with collagen with the biofilm grown without collagen (no collagenase treatment in either case) using the student T-test. Red statistical markers indicate the results of the student T-test comparison between the same collagen concentration, with and without collagenase treatment. Statistically significant differences are shown by * for p < 0.05, ** for p < 0.01, and *** for p < 0.001. The p-values for all tests are shown in Table 5.

**Table 5.**
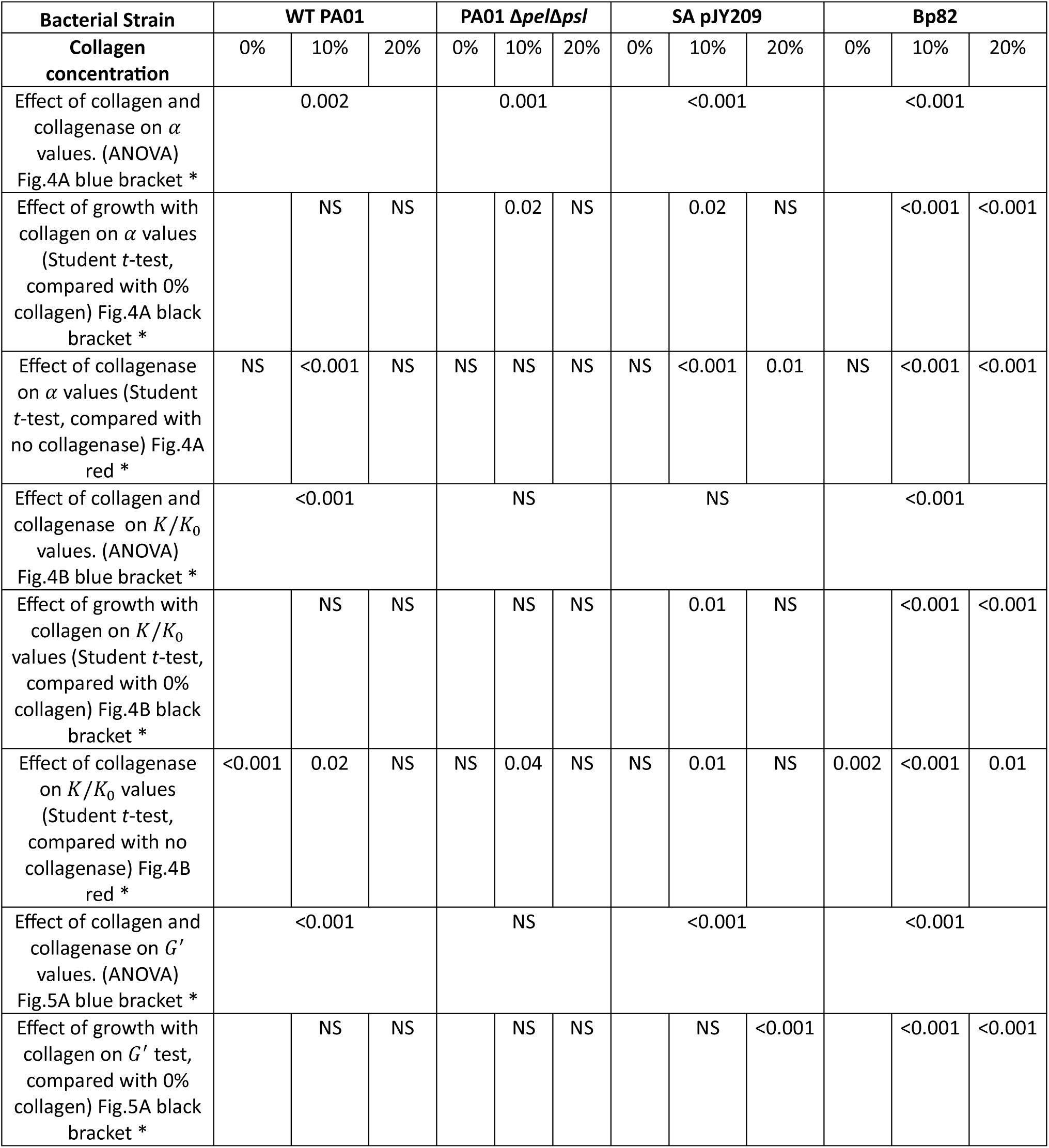
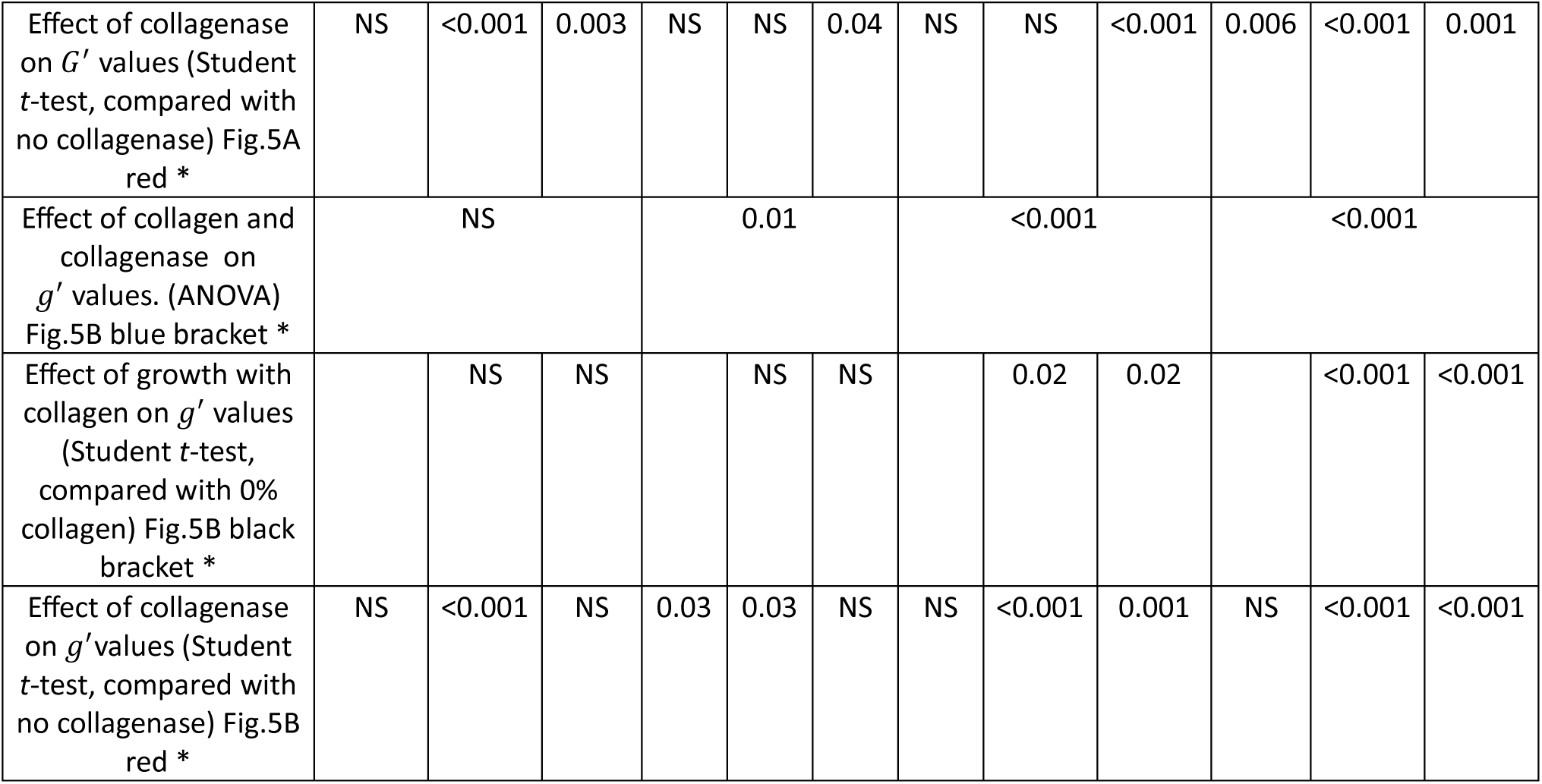
p-values for statistical tests to assess the impact, on microrheology, of growth with collagen and treatment with collagenase. The type of test used (ANOVA or Student t-test) is indicated for each row.

### Collagen enhances microscale elasticity of biofilms that is reduced when treated with collagenase

To further explore the connection between hindered transport, elasticity and phagocytosis, we extract from the MSDs the linear elastic modulus, *G*′ and viscous modulus *G*″, which are measures of how elastic or solid-like and viscous or fluid-like a material is, respectively (see Methods). As shown in **Fig 5A**, we find that growth in the presence of collagen generally increased biofilm elastic modulus, consistent with our previous measurements. The increase is most pronounced for *B. pseudomallei*, in which *G*′ increases over two orders of magnitude from 0% to 20% collagen (∼2 Pa to >200 Pa). Moreover, we find that collagenase treatment decreases *G*′, reaching values comparable to 0% collagen for some strains. We find similar behavior for *G*″ (**SI Fig S13** but the magnitudes of *G*″ are lower than *G*′and the impacts of collagen and collagenase are not as dramatic as for *G*′. These results indicate biofilms are predominantly elastic and that collagen enhances this elasticity. To quantify these effects, we evaluate the relative elasticity *g*′ = *G*′/*G*^*^which is 0 and 1 for a purely viscous and elastic material respectively (**Fig 5B**). We find that for all biofilms and treatments, *g*′ > 0.85, indicating that indeed biofilms are predominantly elastic, in line with our previous bulk rheology measurements [31, 81]. Moreover, we find that growth in the presence of collagen increases *g*′, and enzymatic treatment reduces *g*′.

**Figure 5.**
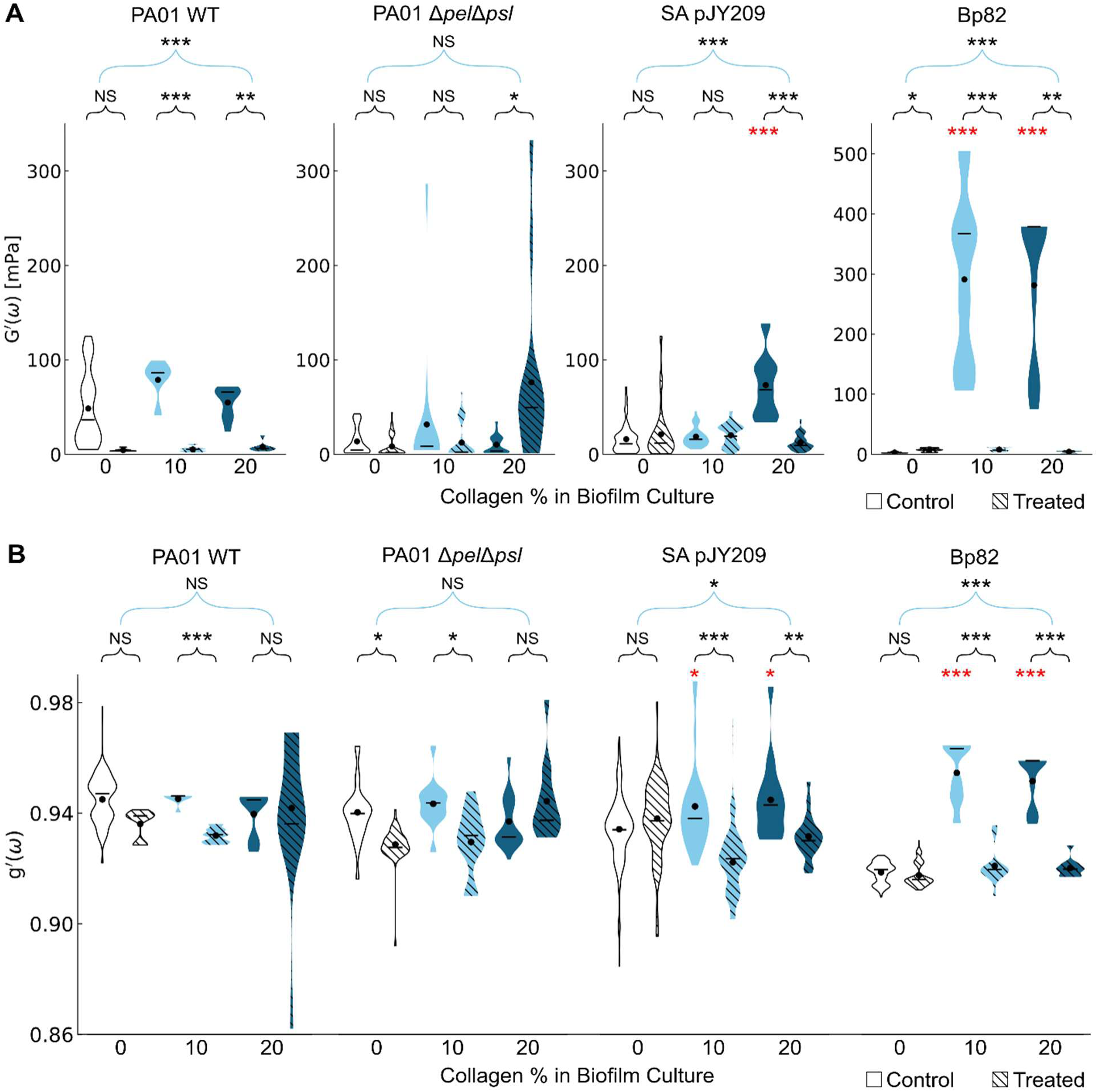
Linear microrheology of biofilms with and without the presence of collagenase. (A) Elastic modulus *G*′ for untreated biofilms grown with different concentrations of collagen (solid violin plots) paired with biofilms that were grown with collagen and then treated with collagenase (hashed violin plots) where the black dot is the weighted average, and the line is the median. (B) The relative elasticity *g*′ = *G*′/*G*^*^ for the untreated and treated biofilm cases. (A and B) Black statistical markers with blue brackets indicate results of the comparison of the six sets of values (all three collagen concentrations, without and with collagenase treatment) using ANOVA. Black statistical markers with black brackets indicate the results of the comparison of the biofilm grown with collagen with the biofilm grown without collagen (no collagenase treatment in either case) using the student T-test. Red statistical markers indicate the results of the student T-test comparison between the same collagen concentration, with and without collagenase treatment. Statistically significant differences are shown by * for p < 0.05, ** for p < 0.01, and *** for p < 0.001. The p-values for all tests are shown in Table 5.

We note that biofilms for microrheology experiments were grown under different conditions than those for engulfment experiments, as described in Methods, and we expect biofilm properties to be dependent on growth conditions [68]. Therefore, we do not expect quantitative agreement between the moduli in both experiments. However, we do expect that the impact on viscoelastic elastic moduli of collagen and collagenase may be similar for both types of growth conditions. That would correspond to an increase by up to two orders of magnitude when biofilms are grown with collagen, and a decrease by up to two orders of magnitude when biofilms are grown with collagenase.

### The correlation of mechanics and phagocytosis

To investigate the relationship between the change in biofilm mechanics and vulnerability to host immune cells, we plotted the normalized change of phagocytic success versus the change in the mechanical property of interest. For each dot in the plots, the vertical (y) coordinate indicates the normalized change in phagocytic success rate (indicated by solid lines in **Figures 2B** and **3B**) and the horizontal (x) coordinate indicates the change in an average mechanical property (indicated by dots in the violin plots in **Figures 4** and **5**).

For three of the four species investigated, the change in normalized phagocytic success rate is positively correlated with the change in anomalous scaling coefficient α (corresponding with a more-fluid transport environment) (**Fig. S14**). For two of these species, there is also a strong positive correlation with an increase in normalized transport K/K_0_ (corresponding with faster motion) (**Fig. S15**).

For at least one of the species investigated, there is a negative correlation between the change in elastic modulus G’ and the change in normalized phagocytic success rate (two others may also show a negative correlation, but the range of ΔG’ values measured make this impossible to conclude with confidence) (**Fig. S16**). For two of the species studied, there is a negative correlation between the change in relative elasticity Δg’ and the change in phagocytic success (**Fig. 17**).

Overall, these findings are consistent with our previous studies showing increased elasticity of *P. aeruginosa* biofilms when grown in collagen [17, 18]. These findings are partly consistent with the idea of a correlation between bulk elasticity of the biofilm (or other larger-than-neutrophil structure) and the ability of neutrophils to phagocytose smaller-than-neutrophils bacteria or beads out of the biofilm or large structure [94, 95]. However, the correlations discussed above are not universal across all strains studied, which suggests that the presence of collagen may also help protect the biofilms in ways other than changing the biofilm viscoelasticity. One possibility is that collagen may alter the fracture mechanics of biofilms, which our current data do not allow us to evaluate. Another possibility is that incorporated collagen, and its degradation by collagenase, may alter the microstructure of biofilms and thereby alter the accessibility of biofilm bacteria to phagocytosing neutrophils, which we briefly survey below.

### Scanning Electron Microscopy uncovers unique microstructures in biofilms grown with collagen

In addition to changes in viscoelasticity, changes in the microstructure of biofilms also have the potential to alter the phagocytic success of attacking neutrophils. For an initial assessment of how collagen and collagenase treatment might impact the microstructure of biofilms, we used our recently published fixation method [96] to preserve the biofilm structure for scanning electron microscopy imaging. We are surprised to observe that a “shield”-like structure is observed at the top of the biofilm for every strain grown with 20% collagen (**Fig. 6**). This “shield”-like crust is not seen on biofilms grown without collagen, and it almost entirely disappears when biofilms are treated with collagenase. Furthermore, compared with biofilms grown without collagen or grown with collagen but then treated with collagenase, biofilms grown with collagen and NOT treated with collagenase had a structure that was better preserved through the fixation process and more bacteria remained attached to the substrate. One possible explanation is that collagen networks in biofilms improved the structural stability of the biofilm and protected the crust from removal during the washing and fixation process. It is also possible that the crust contains collagen as a major structural component and strengthening agent.

**Fig 6.**
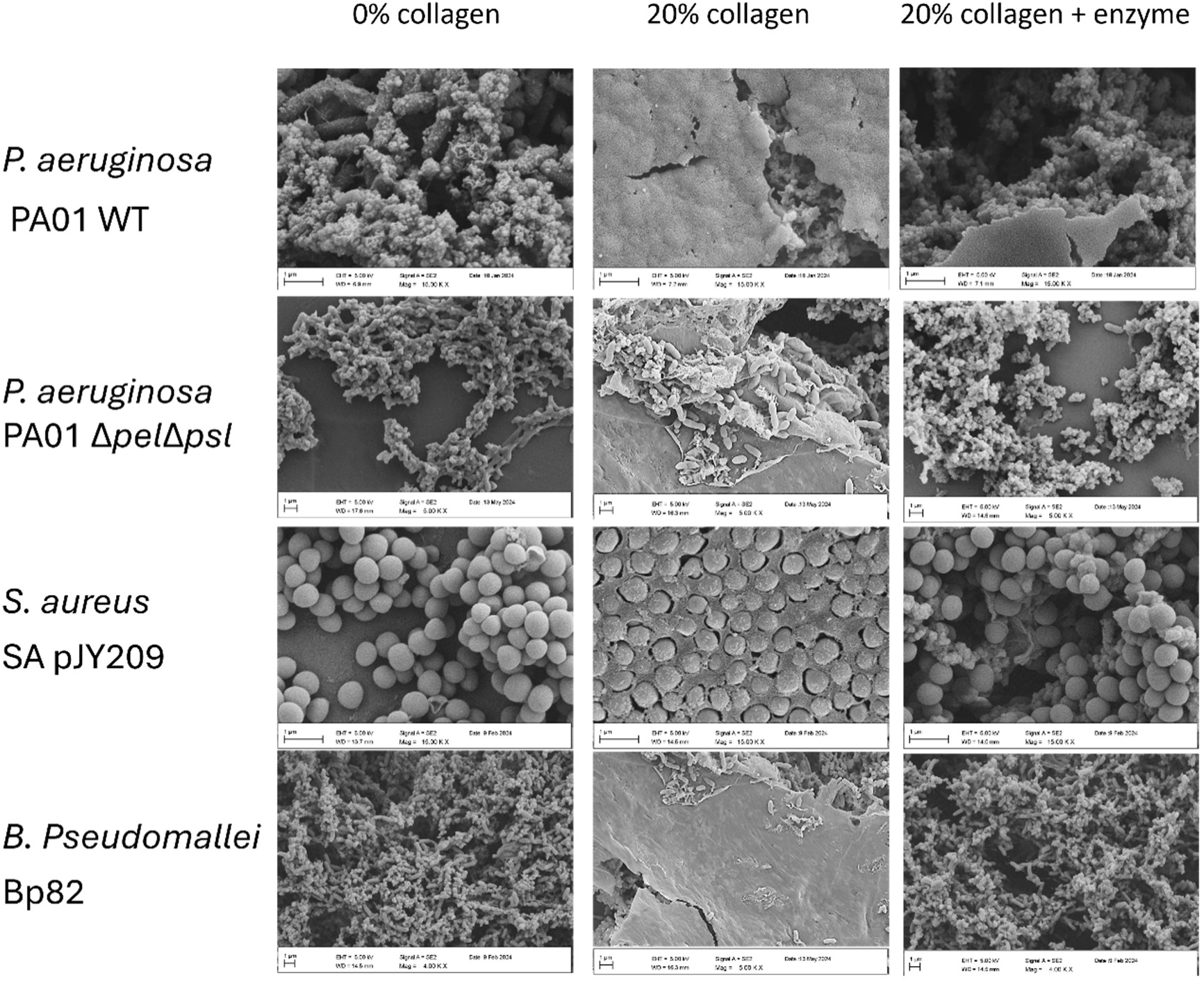
Scanning Electron Microscopy imaging of biofilms grown without collagen (left), with collagen (middle), and grown with collagen then treated with collagenase (right). Scale bars (bottom left of each subpanel) represent 2 μm.

Collagenase treatment may also change the host-bacteria interaction between the biofilms and neutrophils. To visualize this, we used SEM to capture the interaction between PA01 biofilms and human neutrophils. The PA01 biofilms are collected, washed, and incubated, then fixed for SEM imaging. After *P. aeruginosa* biofilms grown with collagen are treated with collagenase, more neutrophils remain attached to the biofilms, which gives them more access to engulf bacteria (**Fig. 7**).

**Figure 7.**
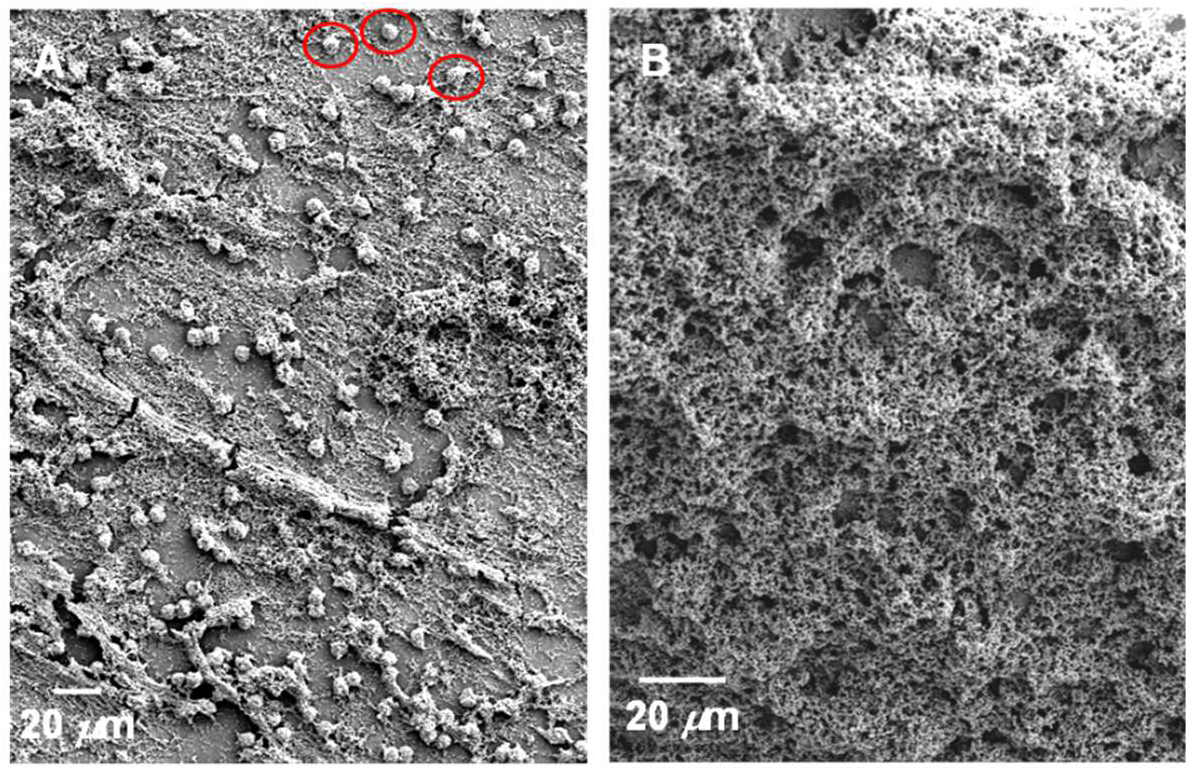
Scanning Electron Microscopy imaging of PA01 WT biofilms grown with collagen that were incubated with neutrophils. The biofilm shown in (A) was treated with collagenase before incubation with neutrophils; the biofilm shown in (B) was not treated with collagenase. Three of the neutrophils in (A) are indicated with red circles. Scale bars are 20 μm.

## Conclusion

Bacterial biofilms are notorious for their ability to evade antibiotics and clearance by the immune system. The elastic response of biofilms allows them to tolerate mechanical stress, while the viscous response allow them to flow along surfaces in response to shear stresses [97, 98]. Biofilms commonly cause persistent infections in many scenarios, including chronic wounds, cystic fibrosis lungs, periodontitis, urinary tract infections (UTIs), and recurrent ear, sinus, and bladder infections [23–28]. One common thing among these infection sites is the abundance of collagen in the connective tissues. Therefore, the ability of biofilms to incorporate host material, such as collagen in the wound bed, may be critically important to researchers developing methods to clear the infection. Here, we present an *in vitro* study of biofilms grown from three highly clinically relevant bacterial species, *P. aeruginosa, S. aureus, and B. pseudomallei*. We find that the incorporation of collagen into the biofilm matrix leads to decreased success of neutrophils at engulfing bacteria, and to increased elasticity. As a corollary, we find that treatment of collagen-supplemented biofilms with collagenase results in increased phagocytic success, and to increased viscosity. This phenomenon was observed across different bacterial strains and species, suggesting a generalized mechanism of host material integration that enhances biofilm resilience.

In conclusion, our study suggests that the incorporation of collagen, and perhaps other host materials, into biofilm infections may contribute to biofilms’ notorious evasion of immune clearance. The presence of host materials in biofilms could potentially provide alternative routes or targets for dispersing the biofilms and enhance the efficacy of the innate immune system at clearing such biofilm infections. By targeting the contributions of host proteins, it may be possible to disrupt biofilm stability and enhance immune clearance in a generalized way, therefore reducing the need for antibiotic treatment and mitigating the development of antibiotic resistance. Such an approach might be beneficial both in cases where the infecting species is known and also in cases wherein biofilm components are not readily known, such as multispecies infections or infections by unknown species.

## Acknowledgements

This work was supported by grants from the National Science Foundation (NSF) (727544 and 2150878, BMMB, CMMI) and the National Institutes of Health (NIH) (1R01AI121500-01A1, NIAID), all to Vernita Gordon, and by a grant from the Air Force Office of Scientific Research (AFOSR) (FA9550-21-1-0361) to Rae Robertson-Anderson. XZ was partly supported by funds from the Trull Centennial Professorship in Physics, of which Vernita Gordon is a fellow. MW was partly supported by the Provost’s Graduate Excellence Fellowship at the University of Texas at Austin. We thank the staff at UT Austin: University Health Service for phlebotomy blood draws, and Center for Biomedical Research Support, especially Michelle Mikesh, for assisting with our SEM imaging. Imaging was done through the Microscopy and Flow Cytometry Facility (RRID:SCR_021756) which is overseen by the Center for Biomedical Research Support at the University of Texas at Austin.

## Notes

### Competing Interest Statement

The authors have declared no competing interest.

### Summary of Updates

Figure 2 was mistakenly repeated as Figure 3 in the previous version. We have now corrected this to show the proper Figure 3.

## References

1. Stewart, P.S., Biophysics of biofilm infection. Pathog Dis, 2014. 70(3): p. 212–8.

2. Stewart, P.S., Mechanisms of antibiotic resistance in bacterial biofilms. Int J Med Microbiol, 2002. 292(2): p. 107–13.

3. Stewart, P.S. and J.W. Costerton, Antibiotic resistance of bacteria in biofilms. Lancet, 2001. 358(9276): p. 135–8.

4. Flemming, H.C., et al., The biofilm matrix: multitasking in a shared space. Nat Rev Microbiol, 2022.

5. Flemming, H.C., T.R. Neu, and D.J. Wozniak, The EPS matrix: the “house of biofilm cells”. J Bacteriol, 2007. 189(22): p. 7945–7.

6. Donlan, R.M., Biofilms: microbial life on surfaces. Emerg Infect Dis, 2002. 8(9): p. 881–90.

7. Costerton, J.W., P.S. Stewart, and E.P. Greenberg, Bacterial biofilms: a common cause of persistent infections. Science, 1999. 284(5418): p. 1318–22.

8. Moser, C., et al., Immune Responses to Pseudomonas aeruginosa Biofilm Infections. Front Immunol, 2021. 12: p. 625597.

9. Versey, Z., et al., Biofilm-Innate Immune Interface: Contribution to Chronic Wound Formation. Front Immunol, 2021. 12: p. 648554.

10. Eriksson, E., et al., Chronic wounds: Treatment consensus. Wound Repair and Regeneration, 2022. 30(2): p. 156–171.

11. Serra, R., et al., Chronic wound infections: the role of Pseudomonas aeruginosa and Staphylococcus aureus. Expert Review of Anti-infective Therapy, 2015. 13(5): p. 605–613.

12. Petras, J.K., et al., Locally Acquired Melioidosis Linked to Environment — Mississippi, 2020–2023. New England Journal of Medicine, 2023. 389(25): p. 2355–2362.

13. Meumann, E.M., et al., Burkholderia pseudomallei and melioidosis. Nature Reviews Microbiology, 2024. 22(3): p. 155–169.

14. Birnie, E., J.J. Biemond, and W.J. Wiersinga, Drivers of melioidosis endemicity: epidemiological transition, zoonosis, and climate change. Current Opinion in Infectious Diseases, 2022. 35(3): p. 196–204.

15. Walker, T.S., et al., Enhanced Pseudomonas aeruginosa biofilm development mediated by human neutrophils. Infection and immunity, 2005. 73(6): p. 3693–3701.

16. Ibanez de Aldecoa, A.L., O. Zafra, and J.E. González-Pastor, Mechanisms and regulation of extracellular DNA release and its biological roles in microbial communities. Frontiers in microbiology, 2017. 8: p. 1390.

17. Rahman, M.U., et al., Effect of collagen and EPS components on the viscoelasticity of Pseudomonas aeruginosa biofilms. Soft Matter, 2021. 17(25): p. 6225–6237.

18. Rahman, M.U., et al., Microrheology of Pseudomonas aeruginosa biofilms grown in wound beds. npj Biofilms and Microbiomes, 2022. 8(1): p. 49.

19. Di Lullo, G.A., et al., Mapping the ligand-binding sites and disease-associated mutations on the most abundant protein in the human, type I collagen. Journal of Biological Chemistry, 2002. 277(6): p. 4223–4231.

20. Varma, S., J.P. Orgel, and J.D. Schieber, Nanomechanics of type I collagen. Biophysical journal, 2016. 111(1): p. 50–56.

21. Makareeva, E. and S. Leikin, Collagen structure, folding and function, in Osteogenesis Imperfecta. 2014, Elsevier. p. 71–84.

22. Chen, Y.-Q., et al., Fibroblast promotes head and neck squamous cell carcinoma cell invasion through mechanical barriers in 3D collagen microenvironments. ACS Applied Bio Materials, 2020. 3(9): p. 6419–6429.

23. Vaca, D.J., et al., Interaction with the host: the role of fibronectin and extracellular matrix proteins in the adhesion of Gram-negative bacteria. Med Microbiol Immunol, 2020. 209(3): p. 277–299.

24. Harsha, L. and M. Brundha, Role of collagen in wound healing. Drug Invention Today, 2020. 13(1).

25. Kaku, M. and M. Yamauchi, Mechano-regulation of collagen biosynthesis in periodontal ligament. J Prosthodont Res, 2014. 58(4): p. 193–207.

26. Wiltberger, G., et al., Mid-and long-term results after replacement of infected peripheral vascular prosthetic grafts with biosynthetic collagen prosthesis. J Cardiovasc Surg (Torino), 2014. 55(5): p. 693–698.

27. Stickler, D.J., Bacterial biofilms in patients with indwelling urinary catheters. Nature clinical practice urology, 2008. 5(11): p. 598–608.

28. Fisher, R.A., B. Gollan, and S. Helaine, Persistent bacterial infections and persister cells. Nature Reviews Microbiology, 2017. 15(8): p. 453–464.

29. Davis-Fields, M., et al., Assaying How Phagocytic Success Depends on the Elasticity of a Large Target Structure. Biophys J, 2019. 117(8): p. 1496–1507.

30. Bakhtiari, L.A., M.J. Wells, and V.D. Gordon, High-throughput assays show the timescale for phagocytic success depends on the target toughness. Biophys Rev (Melville), 2021. 2(3): p. 031402.

31. Kovach, K.N., et al., Specific disruption of established Pseudomonas aeruginosa biofilms using polymer-attacking enzymes. Langmuir, 2020. 36(6): p. 1585–1595.

32. Wells, M., et al., Perspective: The viscoelastic properties of biofilm infections and mechanical interactions with phagocytic immune cells. Front Cell Infect Microbiol, 2023. 13: p. 1102199.

33. Nagoba, B., et al., Treatment of skin and soft tissue infections caused by Pseudomonas aeruginosa—A review of our experiences with citric acid over the past 20 years. Wound Medicine, 2017. 19: p. 5–9.

34. Daum, R.S., Skin and soft-tissue infections caused by methicillin-resistant Staphylococcus aureus. New England Journal of Medicine, 2007. 357(4): p. 380–390.

35. Chewapreecha, C., et al., Global and regional dissemination and evolution of Burkholderia pseudomallei. Nature microbiology, 2017. 2(4): p. 1–8.

36. Nelson, M., et al., The lymphatic system as a potential mechanism of spread of melioidosis following ingestion of Burkholderia pseudomallei. PLoS Neglected Tropical Diseases, 2021. 15(2): p. e0009016.

37. Mitik-Dineva, N., et al., Escherichia coli, Pseudomonas aeruginosa, and Staphylococcus aureus attachment patterns on glass surfaces with nanoscale roughness. Current microbiology, 2009. 58: p. 268–273.

38. Rhodes, K.A. and H.P. Schweizer, Antibiotic resistance in Burkholderia species. Drug Resistance Updates, 2016. 28: p. 82–90.

39. Shockman, G., et al., Bacterial walls, peptidoglycan hydrolases, autolysins, and autolysis. Microbial drug resistance, 1996. 2(1): p. 95–98.

40. Hong, Y. and D.G. Brown, Electrostatic behavior of the charge-regulated bacterial cell surface. Langmuir, 2008. 24(9): p. 5003–5009.

41. Kranjec, C., et al., Staphylococcal Biofilms: Challenges and Novel Therapeutic Perspectives. Antibiotics, 2021. 10(2): p. 131.

42. Izano, E.A., et al., Differential Roles of Poly-N-Acetylglucosamine Surface Polysaccharide and Extracellular DNA in Staphylococcus aureus and Staphylococcus epidermidis Biofilms. Applied and Environmental Microbiology, 2008. 74(2): p. 470–476.

43. Nyanasegran, P.K., et al., Biofilm Signaling, Composition and Regulation in Burkholderia pseudomallei. Journal of Microbiology and Biotechnology, 2023. 33(1): p. 15–27.

44. Wei, Q. and L.Z. Ma, Biofilm Matrix and Its Regulation in Pseudomonas aeruginosa. International Journal of Molecular Sciences, 2013. 14(10): p. 20983–21005.

45. Sadovskaya, I., et al., Structural elucidation of the extracellular and cell-wall teichoic acids of Staphylococcus epidermidis RP62A, a reference biofilm-positive strain. Carbohydrate Research, 2004. 339(8): p. 1467–1473.

46. Tuomanen, E.I., et al., Key Role of Teichoic Acid Net Charge in Staphylococcus aureus Colonization of Artificial Surfaces. Infection and Immunity, 2001. 69(5): p. 3423–3426.

47. Ramsey, D.M. and D.J. Wozniak, Understanding the control of Pseudomonas aeruginosa alginate synthesis and the prospects for management of chronic infections in cystic fibrosis. Molecular Microbiology, 2005. 56(2): p. 309–322.

48. Franklin, M.J., et al., Biosynthesis of the Pseudomonas aeruginosa Extracellular Polysaccharides, Alginate, Pel, and Psl. Frontiers in Microbiology, 2011. 2.

49. Pakkulnan, R., et al., Extracellular DNA facilitates bacterial adhesion during Burkholderia pseudomallei biofilm formation. PLOS ONE, 2019. 14(3): p. e0213288.

50. Foster, T.J., The MSCRAMM Family of Cell-Wall-Anchored Surface Proteins of Gram-Positive Cocci. Trends in Microbiology, 2019. 27(11): p. 927–941.

51. Ymele-Leki, P. and J.M. Ross, Erosion from Staphylococcus aureus biofilms grown under physiologically relevant fluid shear forces yields bacterial cells with reduced avidity to collagen. Applied and environmental microbiology, 2007. 73(6): p. 1834–1841.

52. Grund, M.E., et al., Burkholderia collagen-like protein 8, Bucl8, is a unique outer membrane component of a putative tetrapartite eflux pump in Burkholderia pseudomallei and Burkholderia mallei. Plos one, 2020. 15(11): p. e0242593.

53. Kaewpan, A., et al., Burkholderia pseudomallei pathogenesis in human skin fibroblasts: A Bsa type III secretion system is involved in the invasion, multinucleated giant cell formation, and cellular damage. PLoS One, 2022. 17(2): p. e0261961.

54. David, J., R.E. Bell, and G.C. Clark, Mechanisms of Disease: Host-Pathogen Interactions between Burkholderia Species and Lung Epithelial Cells. Frontiers in Cellular and Infection Microbiology, 2015. 5.

55. McCallon, S.K., D. Weir, and J.C. Lantis, 2nd, Optimizing Wound Bed Preparation With Collagenase Enzymatic Debridement. J Am Coll Clin Wound Spec, 2014. 6(1-2): p. 14–23.

56. Agren, M.S., et al., Collagenase in wound healing: effect of wound age and type. J Invest Dermatol, 1992. 99(6): p. 709–14.

57. Fong, J.N.C. and F.H. Yildiz, Biofilm Matrix Proteins, in Microbial Biofilms. 2015. p. 201–222.

58. Lasa, I. and J.R. Penadés, Bap: A family of surface proteins involved in biofilm formation. Research in Microbiology, 2006. 157(2): p. 99–107.

59. Thi, M.T., D. Wibowo, and B.H.A. Rehm Pseudomonas aeruginosa Biofilms. International Journal of Molecular Sciences, 2020. 21, DOI: 10.3390/ijms21228671.

60. Joo, H.-S. and M. Otto, Molecular Basis of In&#xa0;Vivo Biofilm Formation by Bacterial Pathogens. Chemistry & Biology, 2012. 19(12): p. 1503–1513.

61. Holloway, B.W., Genetic Recombination in Pseudomonas aeruginosa. Microbiology, 1955. 13(3): p. 572–581.

62. Colvin, K.M., et al., The Pel and Psl polysaccharides provide Pseudomonas aeruginosa structural redundancy within the biofilm matrix. Environmental Microbiology, 2012. 14(8): p. 1913–1928.

63. Yarwood, J.M., et al., Quorum Sensing in Staphylococcus aureus Biofilms. Journal of Bacteriology, 2004. 186(6): p. 1838–1850.

64. Propst, K.L., et al., A Burkholderia pseudomallei Δ purM mutant is avirulent in immunocompetent and immunodeficient animals: candidate strain for exclusion from select-agent lists. Infection and immunity, 2010. 78(7): p. 3136–3143.

65. Wells, M.J., H. Currie, and V.D. Gordon, Physiological concentrations of calcium interact with alginate and extracellular DNA in the matrices of Pseudomonas aeruginosa biofilms to impede phagocytosis by neutrophils. Langmuir, 2023. 39(48): p. 17050–17058.

66. Chew, S.C., et al., Dynamic Remodeling of Microbial Biofilms by Functionally Distinct Exopolysaccharides. mBio, 2014. 5(4): p. 10.1128/mbio.01536-14.

67. Chew, S.C., et al., In Situ Mapping of the Mechanical Properties of Biofilms by Particle-tracking Microrheology. Vol. 106. 2015: 1940–087X. e53093.

68. Qi, L. and G.F. Christopher, Rheological variability of Pseudomonas aeruginosa biofilms. Rheologica Acta, 2021. 60(4): p. 219–230.

69. Schindelin, J., et al., Fiji: an open-source platform for biological-image analysis. Nature Methods, 2012. 9(7): p. 676–682.

70. Allan, D., et al., soft-matter/trackpy:v0.6.4 (v0.6.4). Zenodo, 2024.

71. Anderson, S.J., et al., Filament Rigidity Vies with Mesh Size in Determining Anomalous Diffusion in Cytoskeleton. Biomacromolecules, 2019. 20(12): p. 4380–4388.

72. Sokolov, I.M., Models of anomalous diffusion in crowded environments. Soft Matter, 2012. 8(35): p. 9043–9052.

73. Goychuk, I. and T. Pöschel, Fingerprints of viscoelastic subdiffusion in random environments: Revisiting some experimental data and their interpretations. Physical Review E, 2021. 104(3): p. 034125.

74. Anderson, S.J., et al., Subtle changes in crosslinking drive diverse anomalous transport characteristics in actin–microtubule networks. Soft Matter, 2021. 17(16): p. 4375–4385.

75. Robertson-Anderson, R.M., Biopolymer Networks, in Design, dynamics and discovery. 2024, IOP Publishing.

76. McGlynn, J.A., N. Wu, and K.M. Schultz, Multiple particle tracking microrheological characterization: Fundamentals, emerging techniques and applications. Journal of Applied Physics, 2020. 127(20).

77. Squires, T.M. and T.G. Mason, Fluid Mechanics of Microrheology. Annual Review of Fluid Mechanics, 2010. 42(Volume 42, 2010): p. 413–438.

78. Wells, M., M. Mikesh, and V. Gordon, Structure-preserving fixation allows scanning electron microscopy to reveal biofilm microstructure and interactions with immune cells. Journal of Microscopy, 2024. 293(1): p. 59–68.

79. Archer, N.K., et al., Staphylococcus aureus biofilms: properties, regulation, and roles in human disease. Virulence, 2011. 2(5): p. 445–459.

80. Reichhardt, C. and M.R. Parsek, Confocal laser scanning microscopy for analysis of Pseudomonas aeruginosa biofilm architecture and matrix localization. Frontiers in microbiology, 2019. 10: p. 677.

81. Kovach, K., et al., Evolutionary adaptations of biofilms infecting cystic fibrosis lungs promote mechanical toughness by adjusting polysaccharide production. npj Biofilms and Microbiomes, 2017. 3(1): p. 1.

82. Cordova, A., et al., Quantitative morphological analysis of Deinococcus radiodurans elucidates complex dose-dependent nucleoid condensation during recovery from ionizing radiation. Applied and Environmental Microbiology, 2024. 90(7): p. e00108–24.

83. Burtnick Mary, N., J. Brett Paul, and D. DeShazer, Proteomic Analysis of the Burkholderia pseudomallei Type II Secretome Reveals Hydrolytic Enzymes, Novel Proteins, and the Deubiquitinase TssM. Infection and Immunity, 2014. 82(8): p. 3214–3226.

84. Sulik, A. and L. Chyczewski, Immunohistochemical analysis of MMP-9, MMP-2 and TIMP-1, TIMP-2 expression in the central nervous system following infection with viral and bacterial meningitis. Folia Histochemica et Cytobiologica, 2008. 46(4): p. 437–442.

85. Wright, C., et al., Activation of MMP-9 by human lung epithelial cells in response to the cystic fibrosis-associated pathogen Burkholderia cenocepacia reduced wound healing in vitro. American Journal of Physiology-Lung Cellular and Molecular Physiology, 2011. 301(4): p. L575–L586.

86. Dean, R.A., et al., Macrophage-specific metalloelastase (MMP-12) truncates and inactivates ELR+ CXC chemokines and generates CCL2,-7,-8, and-13 antagonists: potential role of the macrophage in terminating polymorphonuclear leukocyte influx. Blood, The Journal of the American Society of Hematology, 2008. 112(8): p. 3455–3464.

87. Wang, L., L. Lankhorst, and R. Bernards, Exploiting senescence for the treatment of cancer. Nature Reviews Cancer, 2022. 22(6): p. 340–355.

88. Di Martino, P., Extracellular polymeric substances, a key element in understanding biofilm phenotype. AIMS microbiology, 2018. 4(2): p. 274.

89. Kosztołowicz, T. and R. Metzler, Diffusion of antibiotics through a biofilm in the presence of diffusion and absorption barriers. Physical Review E, 2020. 102(3): p. 032408.

90. Coppens, B., et al., Anomalous diffusion of nanoparticles in the spatially heterogeneous biofilm environment. iScience, 2023. 26(6).

91. Rozenbaum, R.T., et al., Role of Viscoelasticity in Bacterial Killing by Antimicrobials in Differently Grown. Antimicrob Agents Chemother, 2019. 63(4).

92. Peterson, B.W., et al., Viscoelasticity of biofilms and their recalcitrance to mechanical and chemical challenges. FEMS Microbiol Rev, 2015. 39(2): p. 234–45.

93. Gloag, E.S., et al., Biofilm mechanics: Implications in infection and survival. Biofilm, 2020. 2: p. 100017.

94. Davis-Fields, M., et al., Assaying How Phagocytic Success Depends on the Elasticity of a Large Target Structure. Biophysical Journal, 2019. 117(8): p. 1496–1507.

95. Bakhtiari, L.A., M.J. Wells, and V.D. Gordon, High-throughput assays show the timescale for phagocytic success depends on the target toughness. Biophysics Reviews, 2021. 2(3): p. 031402.

96. Wells, M., M. Mikesh, and V. Gordon, Structure-preserving fixation allows scanning electron microscopy to reveal biofilm microstructure and interactions with immune cells. Journal of Microscopy, 2024. 293(1): p. 59–68.

97. Boudarel, H., et al., Towards standardized mechanical characterization of microbial biofilms: analysis and critical review. NPJ Biofilms Microbiomes, 2018. 4: p. 17.

98. Geisel, S., E. Secchi, and J. Vermant, Experimental challenges in determining the rheological properties of bacterial biofilms. Interface Focus, 2022. 12(6): p. 20220032.

