## Supplementary Material for "Incorporation of collagen into *Pseudomonas aeruginosa, Staphylococcus aureus*, and *Burkholderia pseudomallei* biofilms enhances their elasticity and resistance against phagocytic clearance"

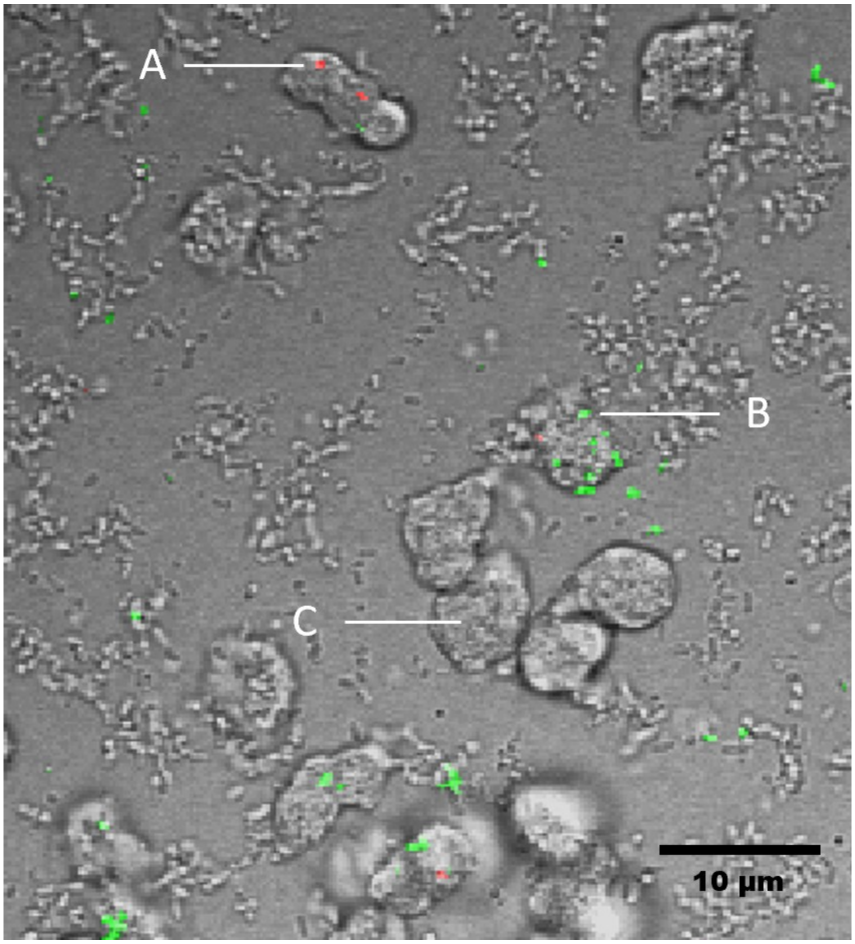


**Fig. S1. Using the pH-dependent stain pH-rodo to distinguish engulfed bacteria from bacteria attached to the neutrophil surface.** A. Bacteria inside the acidic phagosomes emit red fluorescence. B. Bacteria outside of neutrophils only emit green fluorescence (GFP). C. A neutrophil without visible bacteria.

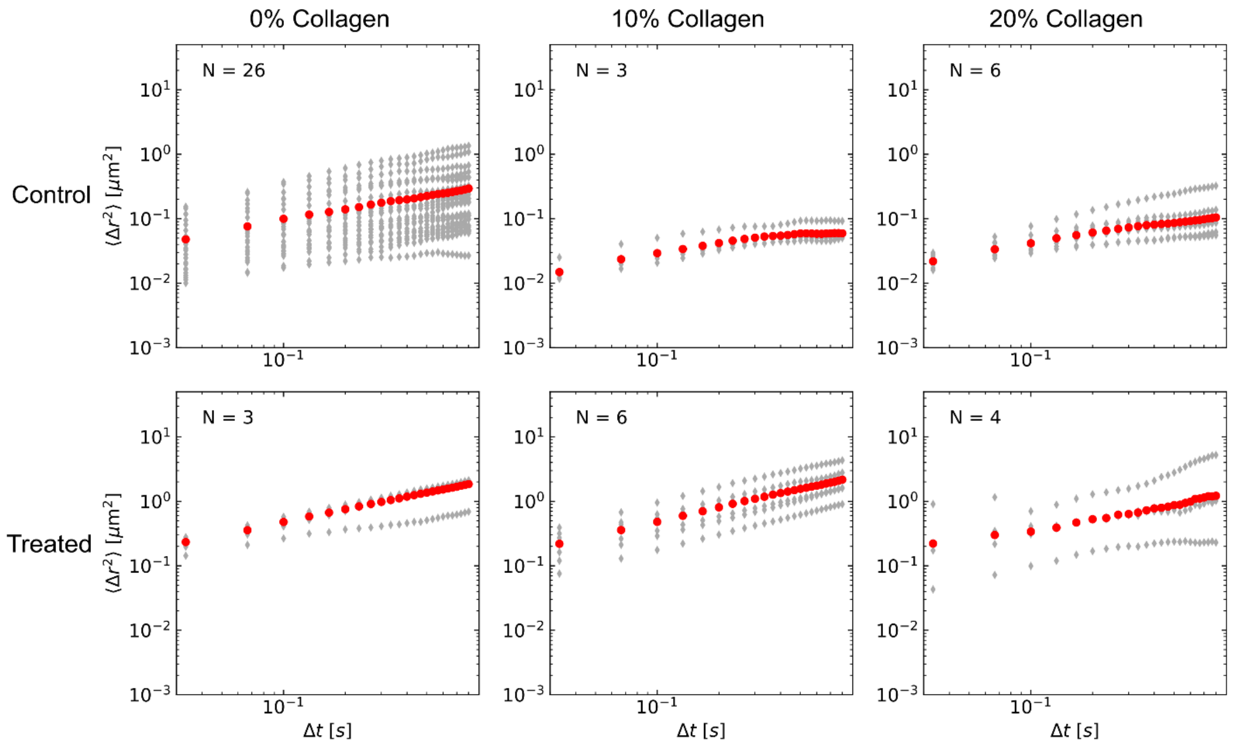


**Fig. S2. Ensemble mean squared displacement (MSD) $\langle \Delta r^2 \rangle$ as a function of lag time Δt for *Pseudomonas aeruginosa* WT biofilms.** Each light grey curve using diamond markers represents one technical replicate the biofilm each condition with N number of trials. The red circle data represents the weighted average MSD-lag time curve where the weight is the number of trajectories divided by the total condition trajectories.

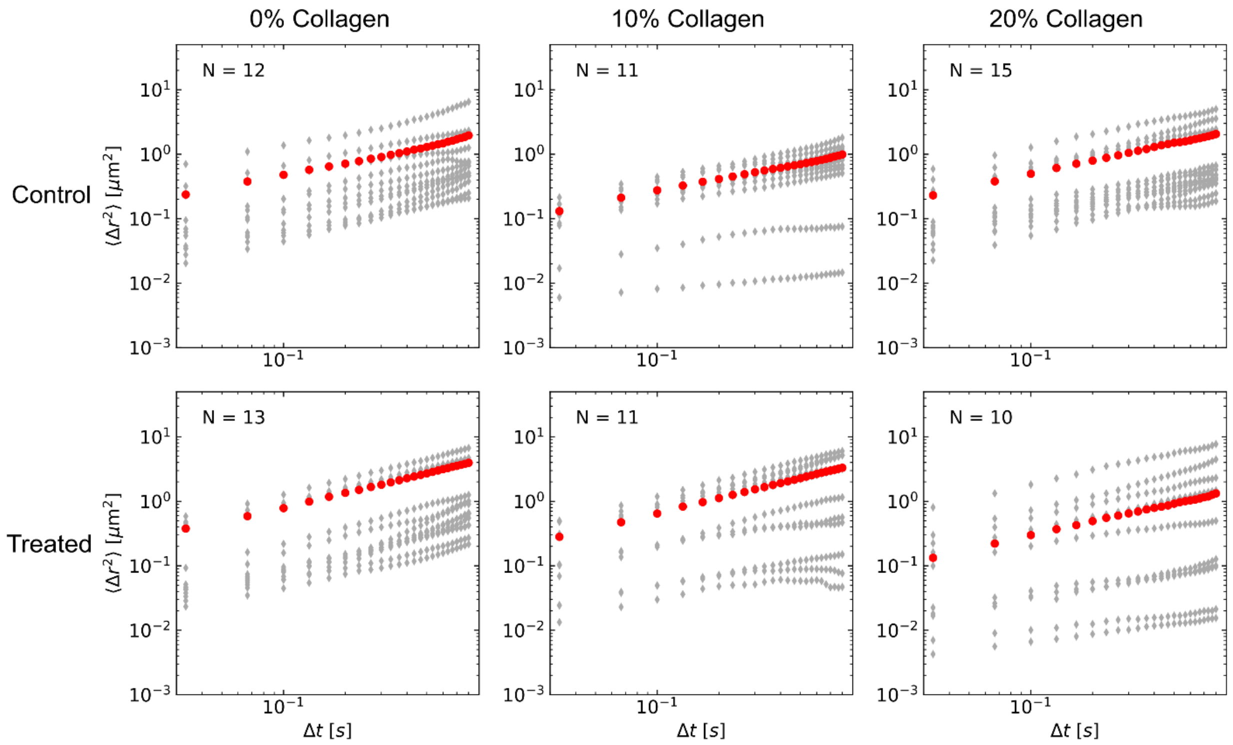


**Fig. S3. Ensemble mean squared displacement (MSD) $\langle \Delta r^2 \rangle$ as a function of lag time Δt for *Pseudomonas aeruginosa* Δ*pel*Δ*psl* biofilms.** Each light grey curve using diamond markers represents one technical replicate the biofilm each condition with N number of trials. The red circle data represents the weighted average MSD-lag time curve where the weight is the number of trajectories divided by the total condition trajectories.

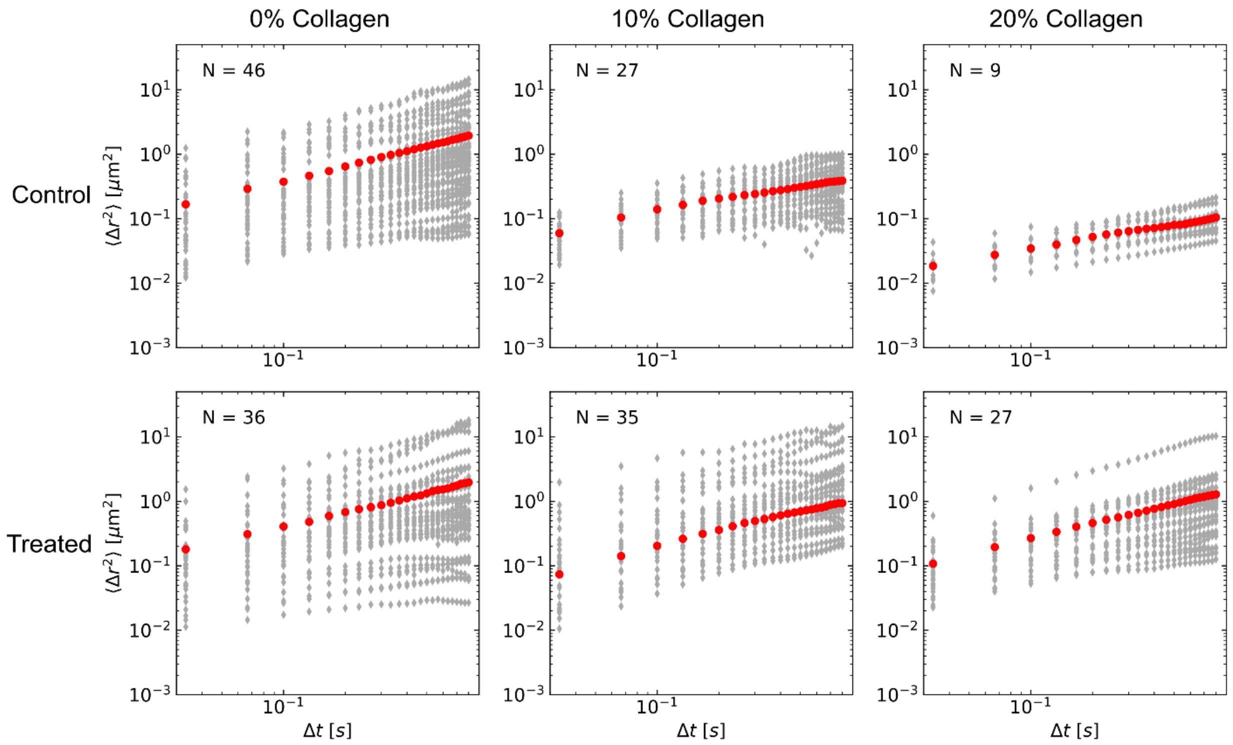


**Fig. S4. Ensemble mean squared displacement (MSD) $\langle \Delta r^2 \rangle$ as a function of lag time Δt for *Staphylococcus aureus* biofilms.** Each light grey curve using diamond markers represents one technical replicate the biofilm each condition with N number of trials. The red circle data represents the weighted average MSD-lag time curve where the weight is the number of trajectories divided by the total condition trajectories.

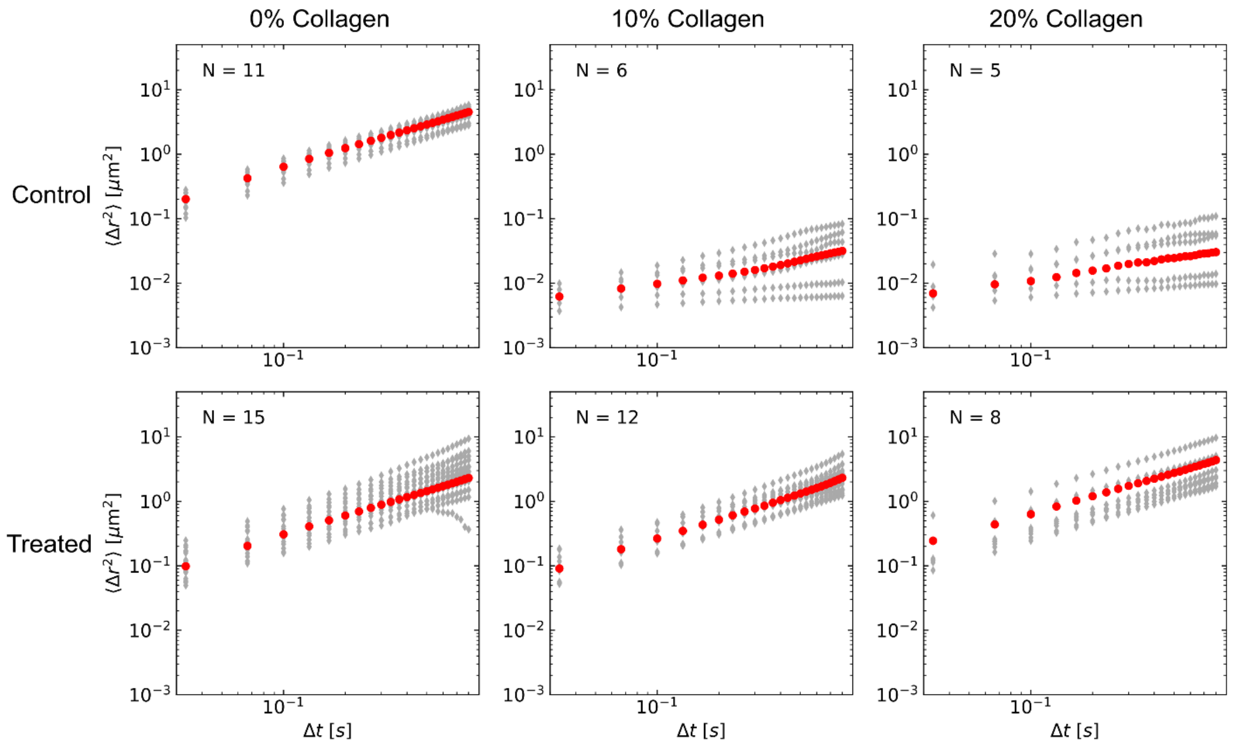


**Fig. S5. Ensemble mean squared displacement (MSD) $\langle \Delta r^2 \rangle$ as a function of lag time Δt for *Burkholderia pseudomallei* biofilms.** Each light grey curve using diamond markers represents one technical replicate the biofilm each condition with N number of trials. The red circle data represents the weighted average MSD-lag time curve where the weight is the number of trajectories divided by the total condition trajectories.

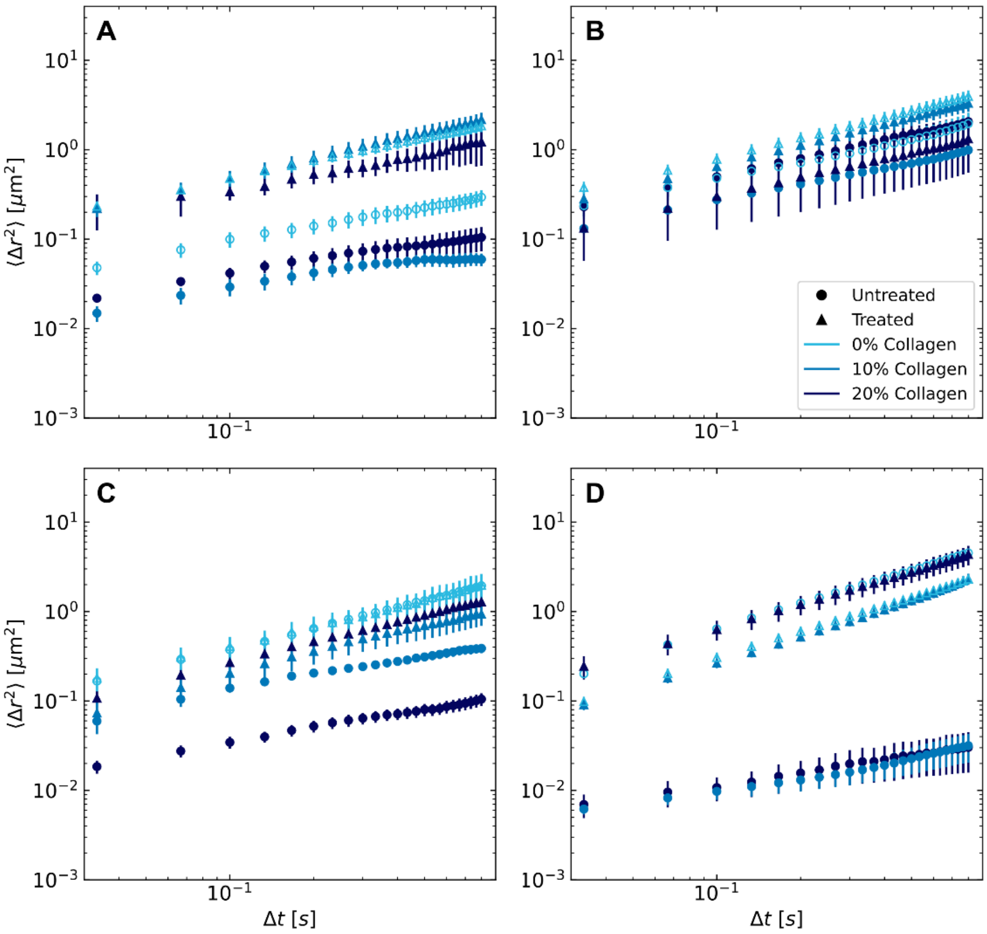


**Fig. S6. Weighted average mean squared displacement (MSD) as a function of lag time Δt for all biofilm growth and treatment conditions:** (A) *Pseudomonas aeruginosa* WT, (B) *Pseudomonas aeruginosa* Δ*pel*Δ*psl*, (C) *Staphylococcus aureus*, and (D) *Burkholderia pseudomallei*.

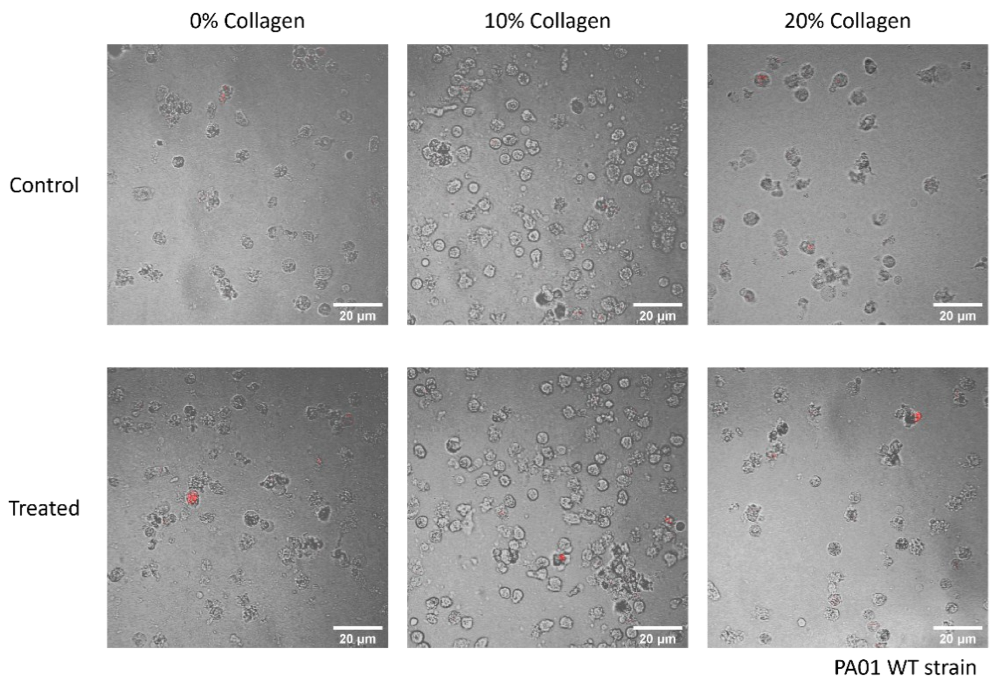


**Fig. S7. Micrographs of neutrophils after they had been incubated with biofilms for 30 minutes, for every condition tested, for PA01 WT biofilms.** Before being incubated with neutrophils, the biofilms were grown with 0%,10%, and 20% collagen, and treated with collagenase or PBS (as control). Scale bars are 20 μm.

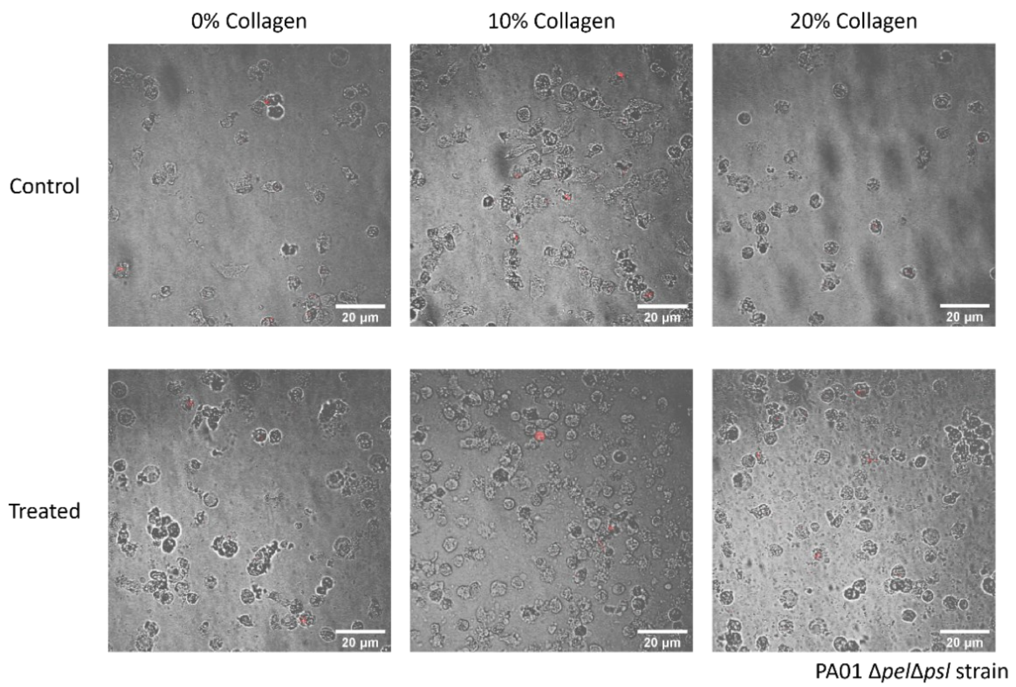


**Fig. S8. Micrographs of neutrophils after they had been incubated with biofilms for 30 minutes, for every condition tested, for PA01 Δ*pel* Δ*psl* biofilms.** Before being incubated with neutrophils, the biofilms were grown with 0%,10%, and 20% collagen, and treated with collagenase or PBS (as control). Scale bars are 20 µm.

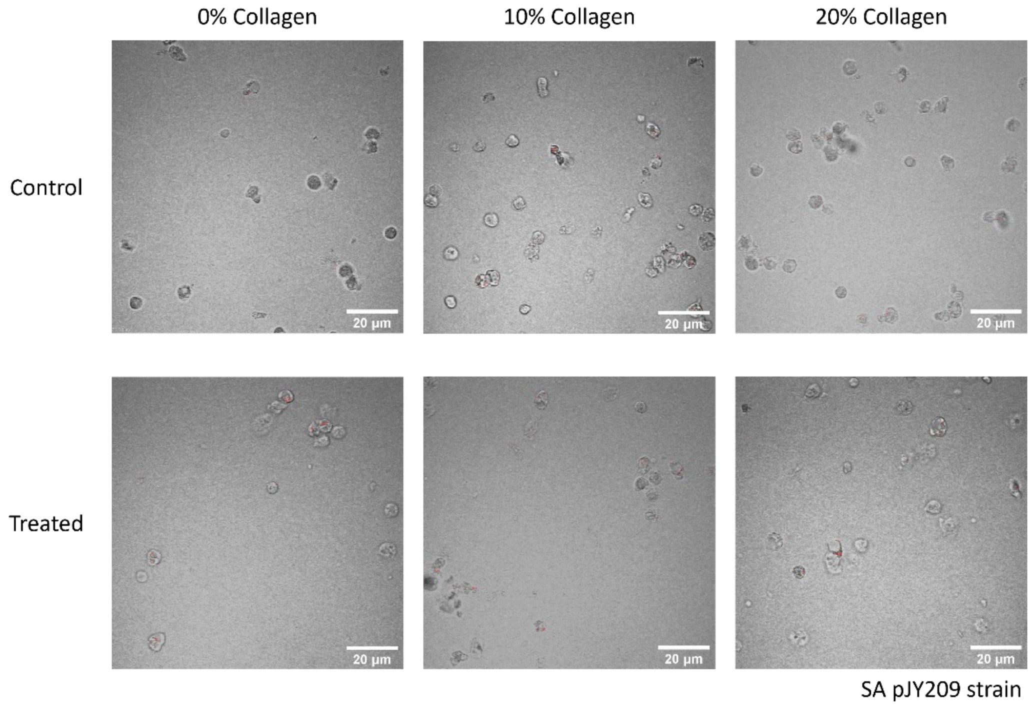


**Fig. S9. Micrographs of neutrophils after they had been incubated with biofilms for 30 minutes, for every condition tested, for *S. aureus* SA pJY209.** Before being incubated with neutrophils, the biofilms were grown with 0%,10%, and 20% collagen, and treated with collagenase or PBS (as control). Scale bars are 20 μm.

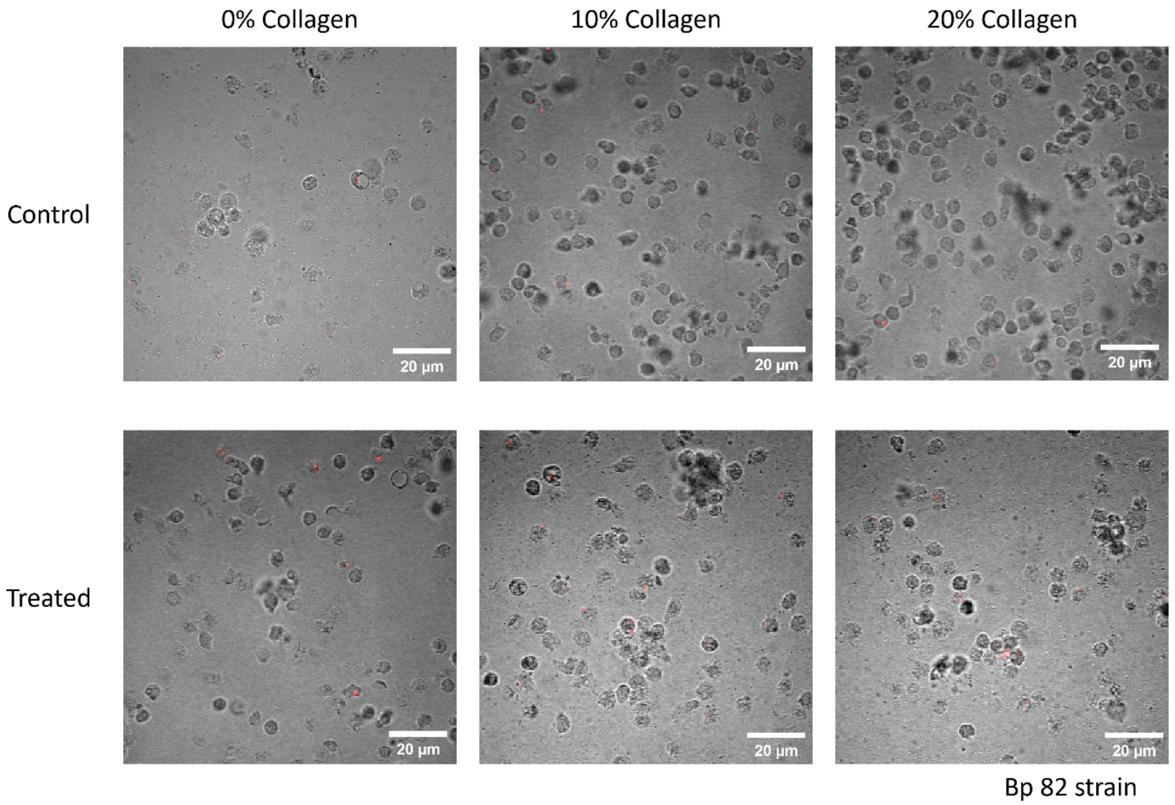


**Fig. S10. Micrographs of neutrophils after they had been incubated with biofilms for 30 minutes, for every condition tested, for *B. pseudomallei* Bp82.** Before being incubated with neutrophils, the biofilms were grown with 0%,10%, and 20% collagen, and treated with collagenase or PBS (as control). Scale bars are 20 µm.

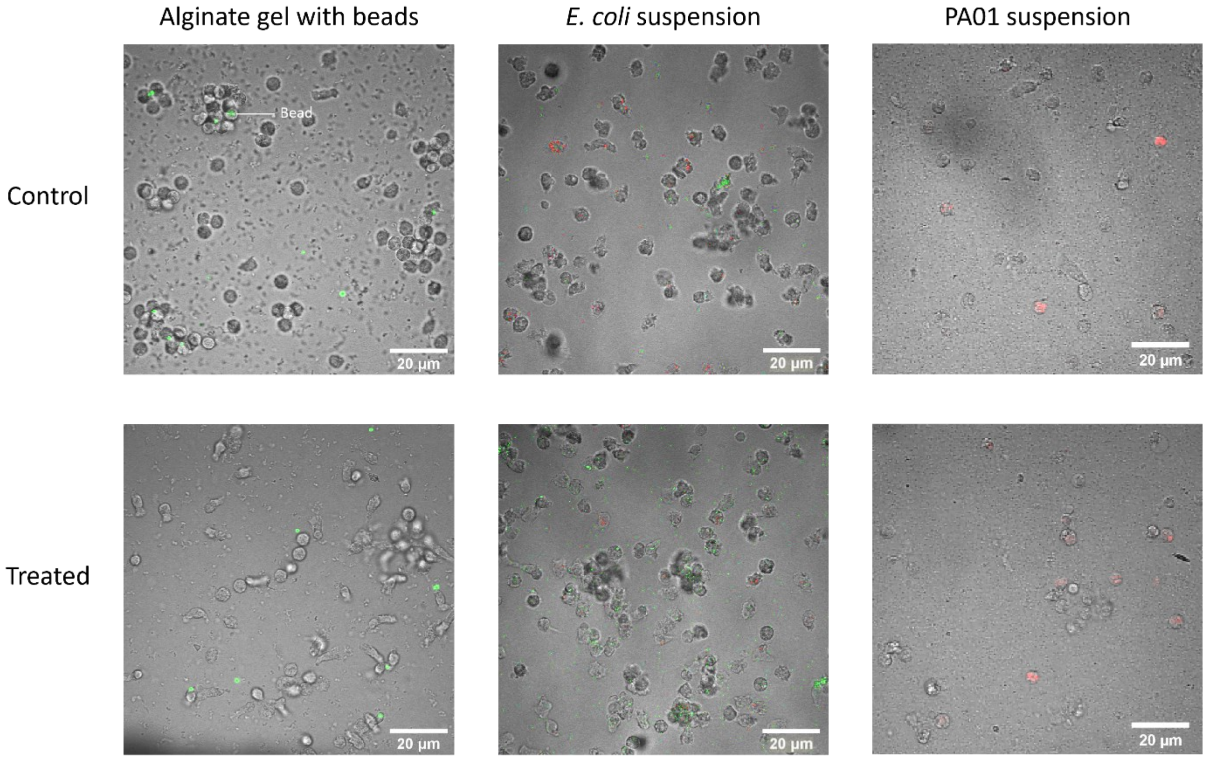


**Fig. S11. Micrographs of neutrophils after they had been incubated with non-biofilm controls – alginate gel containing beads, a suspension of planktonic *Escherichia coli* bacteria, and a suspension of planktonic PA01 P. *aeruginosa* bacteria.** To explore the effect of collagenase on neutrophils, *E. coli* and *P. aeruginosa* suspensions and alginate gel containing tracer beads were treated with collagenase or PBS, then incubated with neutrophils.

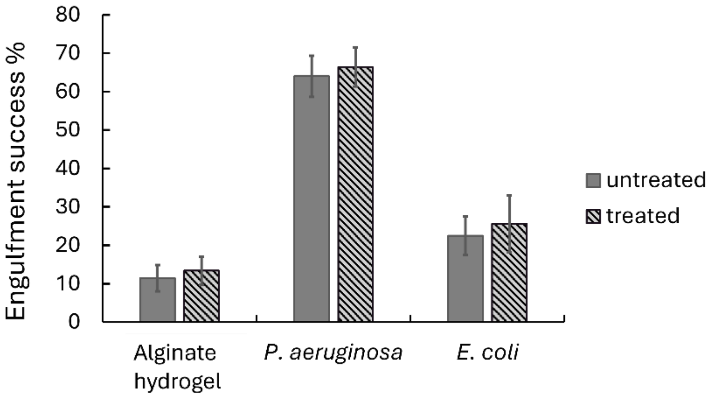


**Fig. S12. Results of control experiments for evaluating the effects of collagenase treatment on neutrophils.** The engulfment assays were done with fluorescent tracer beads initially embedded in alginate hydrogel (left), and with suspensions of *P. aeruginosa* (center) and *E. coli* (right). The engulfment success rate was compared between materials treated with and without collagenase. In both datasets, the student t-test and ANOVA test showed no significance between treated and untreated samples. The p-values and replicate numbers are shown in Table 3. N=3 biological replicates.

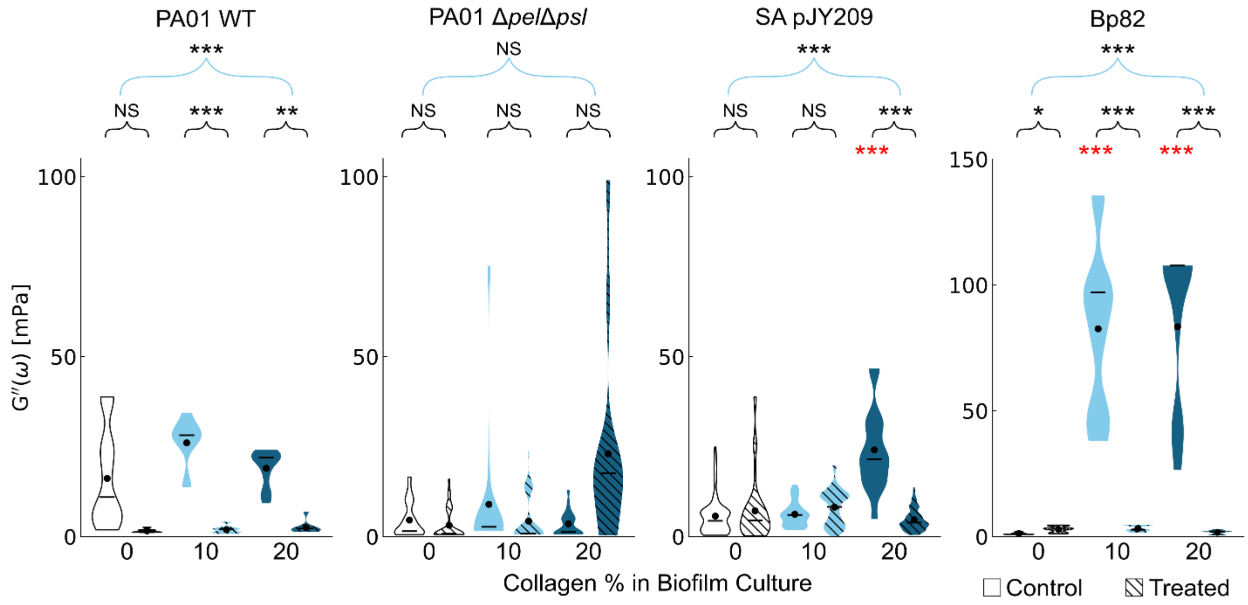


**Fig S13. Linear viscous modulus $G''$ of biofilms with and without the presence of collagenase.** Untreated biofilms (control, solid violin plots) paired with biofilms that were treated with collagenase (hashed violin plots) where the black dot is the weighted average, and the line is the median Black statistical markers with blue brackets indicate results of the comparison of the six sets of values (all three collagen concentrations, without and with collagenase treatment) using ANOVA. Black statistical markers with black brackets indicate the results of the comparison of the biofilm grown with collagen with the biofilm grown without collagen (no collagenase treatment in either case) using the student T-test. Red statistical markers indicate the results of the student T-test comparison between the same collagen concentration, with and without collagenase treatment. Statistically significant differences are shown by * for $p < 0.05$, ** for $p < 0.01$, and *** for $p < 0.001$.

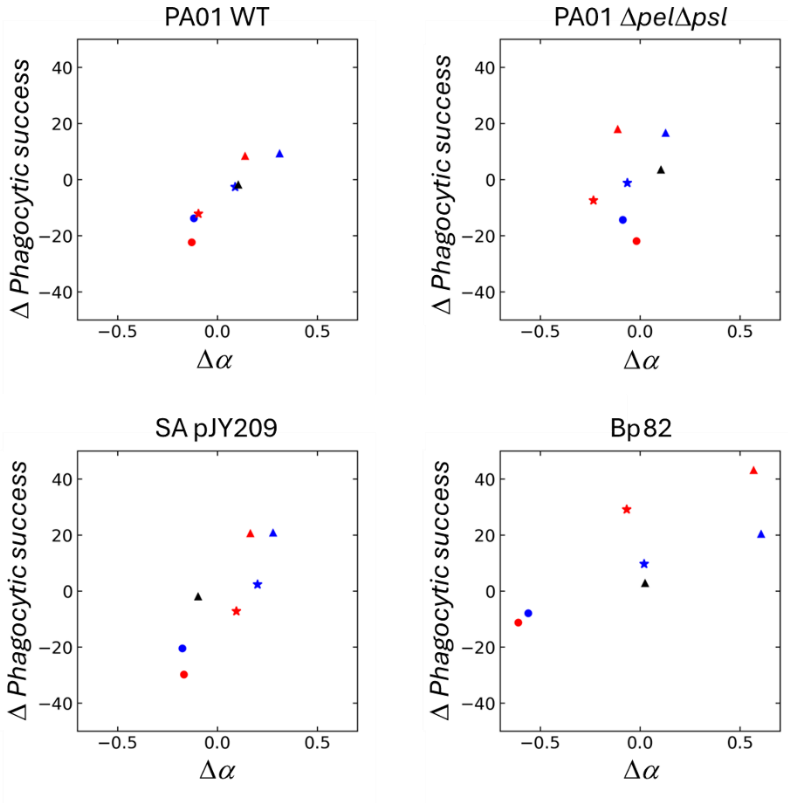


**Fig S14. Scatter plots of the change in phagocytic success versus the associated change in α value.** *P. aeruginosa* (PA01 WT), *S. aureus* (SA pJY209), and *Burkholderia pseudomallei* (Bp82) show a correlation between these changes; the double knockout mutant PA01 Δ*pel*Δ*psl*, which does not make Pel and Psl (the two major matrix polymers made by PA01 biofilms) does not. Colors and shapes of symbols indicate the experimental cases whose averages were used to obtain each value. Circles indicate changes associated with growth with collagen; blue indicates the difference in the value measured for 10% collagen and the value measured for 0% collagen and red indicates the difference in the value measured for 20% collagen and the value measured for 0% collagen. Triangles indicate changes associated with treatment with collagenase; black indicates the difference in the value induced upon collagenase treatment of a 0% collagen case, blue indicates the difference in the value induced upon collagenase treatment of a 10% collagen case, and red, black indicates the difference in the value induced upon collagenase treatment of a 20% collagen case. Stars indicate the residual differences after both compared biofilms were treated with collagenase; blue indicates the 10% collagen case compared with the 0% collagen case, and red indicates the 10% collagen case compared with the 0% collagen case.

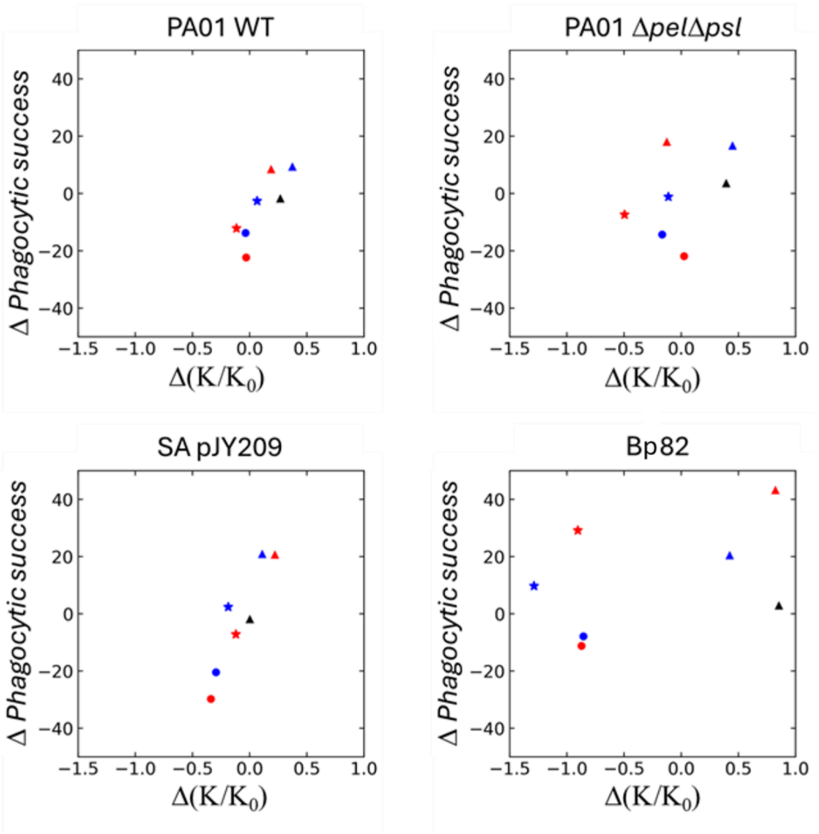


**Fig S15. Scatter plots of the change in phagocytic success versus the associated change in $K/K_0$.** *P. aeruginosa* (PA01 WT) and *S. aureus* (SA pJY209) show a correlation between these changes. If *Burkholderia pseudomallei* (Bp82) shows a correlation at all, it is weak. The double knockout mutant PA01 $\Delta$*pel*$\Delta$*psl*, which does not make Pel and Psl (the two major matrix polymers made by PA01 biofilms) shows no correlation. Colors and shapes of symbols indicate the experimental cases whose averages were used to obtain each value. Circles indicate changes associated with growth with collagen; blue indicates the difference in the value measured for 10% collagen and the value measured for 0% collagen and red indicates the difference in the value measured for 20% collagen and the value measured for 0% collagen. Triangles indicate changes associated with treatment with collagenase; black indicates the difference in the value induced upon collagenase treatment of a 0% collagen case, blue indicates the difference in the value induced upon collagenase treatment of a 10% collagen case, and red, black indicates the difference in the value induced upon collagenase treatment of a 20% collagen case. Stars indicate the residual differences after both compared biofilms were treated with collagenase; blue indicates the 10% collagen case compared with the 0% collagen case, and red indicates the 10% collagen case compared with the 0% collagen case.

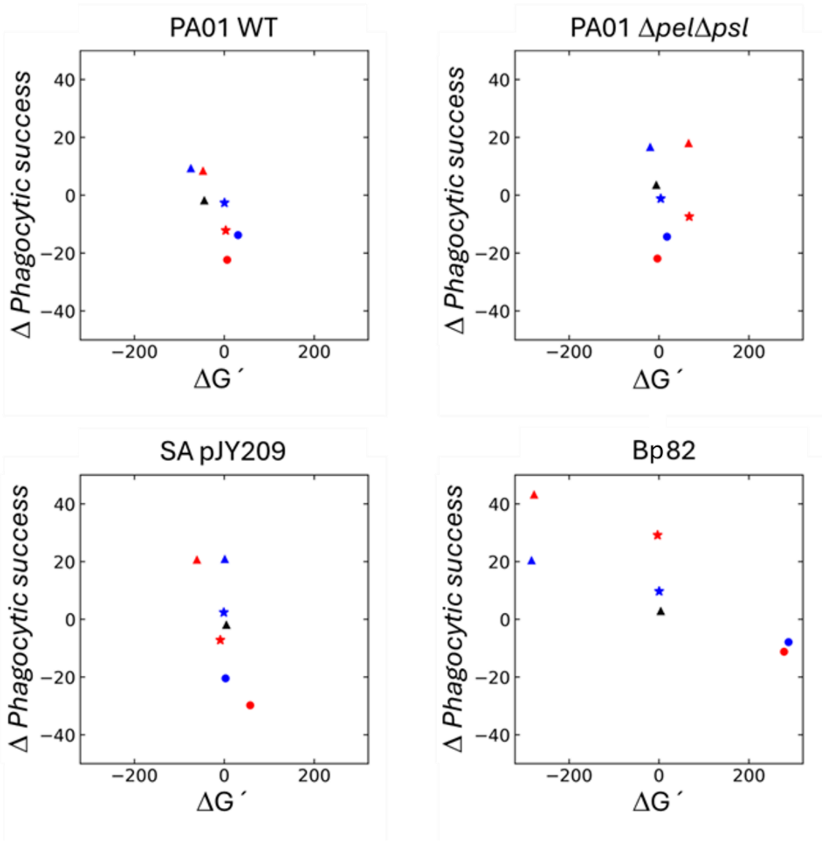


**Fig S16. Scatter plots of the change in phagocytic success versus the associated change in G´.** *Burkholderia pseudomallei* (Bp82) shows a correlation between these changes. *P. aeruginosa* (PA01 WT) and *S. aureus* (SA pJY209) may show a correlation between these changes, but the range of ΔG´ values measured for these biofilms does not allow that conclusion to be confidently drawn. The double knockout mutant PA01 Δ*pel*Δ*psl*, which does not make Pel and Psl (the two major matrix polymers made by PA01 biofilms) shows no correlation. Colors and shapes of symbols indicate the experimental cases whose averages were used to obtain each value. Circles indicate changes associated with growth with collagen; blue indicates the difference in the value measured for 10% collagen and the value measured for 0% collagen and red indicates the difference in the value measured for 20% collagen and the value measured for 0% collagen. Triangles indicate changes associated with treatment with collagenase; black indicates the difference in the value induced upon collagenase treatment of a 0% collagen case, blue indicates the difference in the value induced upon collagenase treatment of a 10% collagen case, and red, black indicates the difference in the value induced upon collagenase treatment of a 20% collagen case. Stars indicate the residual differences after both compared biofilms were treated with collagenase; blue indicates the 10% collagen case compared with the 0% collagen case, and red indicates the 10% collagen case compared with the 0% collagen case.

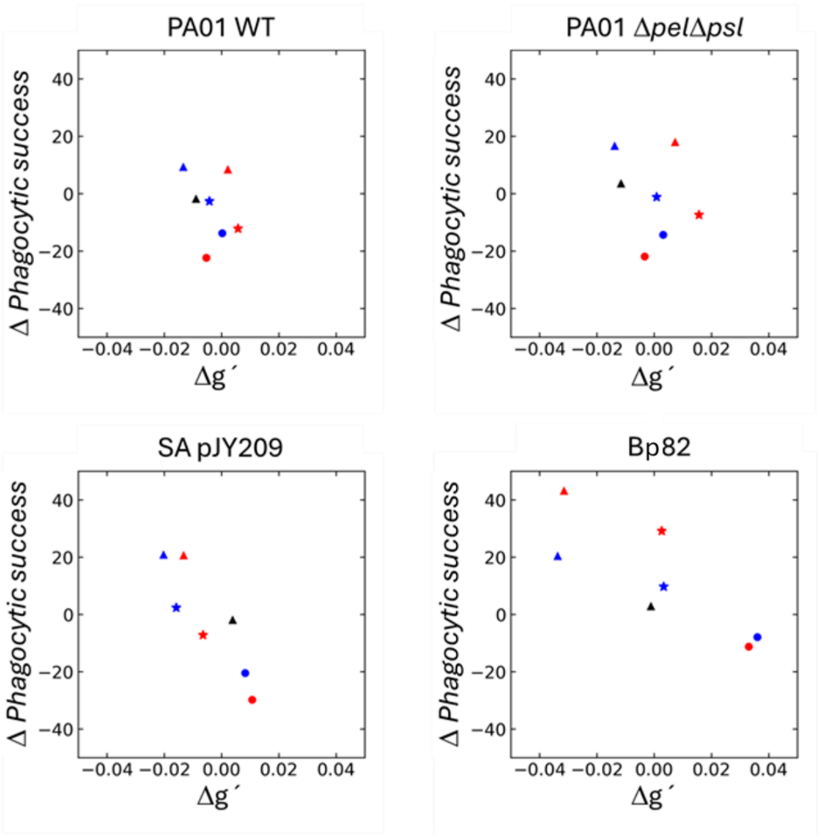


**Fig S17. Scatter plots of the change in phagocytic success versus the associated change in g´.** *S. aureus* (SA pJY209) and *Burkholderia pseudomallei* (Bp82) show a correlation between these changes. The two strains of *P. aeruginosa* studied (PA01 WT and PA01 Δ*pel*Δ*psl*) do not. Colors and shapes of symbols indicate the experimental cases whose averages were used to obtain each value. Circles indicate changes associated with growth with collagen; blue indicates the difference in the value measured for 10% collagen and the value measured for 0% collagen and red indicates the difference in the value measured for 20% collagen and the value measured for 0% collagen. Triangles indicate changes associated with treatment with collagenase; black indicates the difference in the value induced upon collagenase treatment of a 0% collagen case, blue indicates the difference in the value induced upon collagenase treatment of a 10% collagen case, and red, black indicates the difference in the value induced upon collagenase treatment of a 20% collagen case. Stars indicate the residual differences after both compared biofilms were treated with collagenase; blue indicates the 10% collagen case compared with the 0% collagen case, and red indicates the 10% collagen case compared with the 0% collagen case.
